# Pangenome sequence evolution within human gut microbiomes is explained by gene-specific rather than host-specific selective pressures

**DOI:** 10.1101/2020.09.30.319558

**Authors:** Arnaud N’Guessan, Ilana Lauren Brito, Adrian W.R. Serohijos, B. Jesse Shapiro

## Abstract

Pangenomes – the cumulative set of genes encoded by a species – arise from evolutionary forces including horizontal gene transfer (HGT), drift, and selection. The relative importance of drift and selection in shaping pangenome structure has been recently debated, and the role of sequence evolution (point mutations) within mobile genes has been largely ignored, with studies focusing mainly on patterns of gene presence or absence. The effects of drift, selection, and HGT on pangenome evolution likely depends on the time scale being studied, ranging from ancient (*e.g.*, between distantly related species) to recent (*e.g.*, within a single animal host), and the unit of selection being considered (*e.g.*, the gene, whole genome, microbial species, or human host). To shed light on pangenome evolution within microbiomes on relatively recent time scales, we investigate the selective pressures acting on mobile genes using a dataset that previously identified such genes in the gut metagenomes of 176 Fiji islanders. We mapped the metagenomic reads to mobile genes to call single nucleotide variants (SNVs) and calculate population genetic metrics that allowed us to infer deviations from a neutral evolutionary model. We found that mobile gene sequence evolution varied more by gene family than by human social attributes, such as household or village membership, suggesting that selection at the level of gene function is most relevant on these short time scales. Patterns of mobile gene sequence evolution could be qualitatively recapitulated with a simple evolutionary simulation, without the need to invoke an adaptive advantage of mobile genes to their bacterial host genome. This suggests that, at least on short time scales, a majority of the pangenome need not be adaptive. On the other hand, a subset of gene functions including defense mechanisms and secondary metabolism showed an aberrant pattern of molecular evolution, consistent with species-specific selective pressures or negative frequency-dependent selection not seen in prophages, transposons, or other gene categories. That mobile genes of different functions behave so differently suggests stronger selection at the gene level, rather than at the genome level. While pangenomes may be largely adaptive to their bacterial hosts on longer evolution time scales, here we show that, on shorter “human” time scales, drift and gene-specific selection predominate.

## INTRODUCTION

Human gut microbial communities (or microbiomes) impact diverse aspects of human health, such as food digestion, nutritional uptake, immunity, and inflammation^1,2^. The gut microbiome is shaped by both ecological factors, such as shifts in species abundance or strain replacements, and evolutionary forces, such as mutation, horizontal gene transfer (HGT), drift and selection^3^. In particular, microbes in the gut dynamically and frequently exchange genetic material through HGT^4^, resulting in pangenomes (the total set of genes observed in all members of a species or population) which are often much larger than an individual genome size ^5-7^. Some studies have shown that horizontally transferred (mobile) genes could contribute to environmental adaptation, notably through the propagation of antibiotic resistance^5^. However, there are contexts in which pangenome evolution could be driven more by drift than by selection. For instance, the evolution of endosymbionts or intracellular pathogens, which have small effective population sizes, is generally driven by drift, resulting in small pangenomes^8^. In contrast, selection seems to play a bigger role in free-living microbes, like hydrothermal vent bacteria^9^. Whether pangenome evolution is mainly driven by selection (an adaptive model) or drift (a non-adaptive or neutral model) is a question that has generated some controversy ^6,7,10,11^.

Answering this question depends on the time scale being studied. For example, long-term evolution (*e.g.* among distantly related species or among all extant members of a species) versus near-term evolution (*e.g.* among a locally coexisting population of a species) may experience different regimes of drift and selection. On long time scales, using data from distantly related genomes that diverged millions of years ago ^6,7^, and at the whole-genome scale, adaptive and non-adaptive models have been proposed and are still a source of contention. A model in which gene gain by HGT is predominantly adaptive provides a good fit to distantly related genomes from the NCBI database^7^. In that work, Sela and collaborators developed a model of prokaryotic genome size evolution that includes gene gain, gene loss, and their fitness effects ^1^. In their model, gene gain and loss maintain genome size equilibrium and have opposite fitness effects. The model also accounts for species effective population size (*N*_*e*_), which measures genetic diversity and effectiveness of selection in a population, and is dependent on census population size and its fluctuations^11^ as well as on varying intensities of purifying, positive, or fluctuating natural selection. From simulations of this model, they found that a scenario in which gene gain is, on average, slightly beneficial best fits genome size and nucleotide diversity data from 707 prokaryotic genomes. Based on a synthesis of population genomic data and models including Sela and collaborators’ model ^7^, another group led by McInerney and collaborators argued that an adaptive model best explains pangenome evolution because more diverse pangenomes tend to arise in species with larger *N*_*e*_ due to beneficial gene gain, higher efficacy of selection, and a large number of micro-niches available to the species ^6^.

In contrast, Andreani and collaborators observed that genome fluidity, defined as the ratio between the number of unique gene families and the average number of gene families between random genome pairs, significantly correlates with synonymous nucleotide diversity in 90 bacterial species. Although this does not exclude a role for selection, the observation is most parsimoniously explained by a neutral model. Similarly, Bobay and Ochman observed that gene turnover does not significantly correlate with *dN/dS*, which measures selection on protein-coding genes ^11^. They also found that *N*_*e*_ correlates positively with pangenome size for most of the 153 analyzed prokaryotic species. Similar to McInerney and collaborators, they attributed this to an increased effectiveness of selection in species with larger *N*_*e*_ and that most of the accessory genes, those that are present in some but not all strains of a species, are slightly beneficial ^6^. The fact that Bobay and Ochman found evidence for both adaptive and neutral pangenome evolution may seem contradictory ^11^. However, they reconciled these observations by proposing a nearly neutral model of drift-barrier evolution. This model describes the balance between selection and drift. More precisely, it assumes that most accessory genes in the pangenome are slightly beneficial, such that they can be considered neutral when *N*_*e*_ is small, but they can escape the effects of drift and spread when the selective coefficient *s* exceeds 1/*N*_*e*_.

Resolving the balance of evolutionary forces influencing pangenomes also depends on the biological scale or unit of evolution. For example, the consequences of selection at the level of single genes, whole genomes, microbial species or human hosts could yield different patterns. The studies above focused on adaptation at the whole-genome level, but selection also acts at the level of individual genes^9,12,13^. Mobile genes in particular may have their own *N*_*e*_, which could be distinct from the *N*_*e*_ of the whole genome of a species^13^. For example, there is an entire class of mobile genes, including phage and other “selfish” elements that have effectively instantaneous HGT rates^14^. Other mobile genes may provide rapid adaptive value to their bacterial hosts, such as in the gut microbiome of humans with different diets or lifestyles^1^. Therefore, based on their patterns of presence or absence, some mobile genes appear to be selected to favour their own replication (selfish) while others may provide benefits to their bacterial or even human hosts^15^.

All the studies above investigated pangenome evolution among distantly related genomes over relatively ancient time scales. Yet selective pressures might differ on recent and shorter evolutionary time scales, such as within local populations of bacteria over dozens rather than millions of years. However, a targeted investigation of the population genetics of mobile genes on short time scales is still missing. To study pangenome evolution on shorter evolutionary time scales and at the level of individual genes, we used a dataset from Brito and collaborators composed of 37,853 mobile genes involved in recent HGT events in the human gut ^1^. We mapped metagenomic reads from a cohort of 176 Fiji islander gut microbiomes to this set of mobile genes. From the mapped reads, we identified single nucleotide variants (SNVs) segregating within microbiomes, from which we calculated population genetic metrics such as *dN/dS* and *Tajima’s D* that contain information about evolutionary and demographic history of mobile genes. In contrast to studies over longer evolutionary time scales, which have concluded that pangenome evolution is adaptive, we find that many aspects of pangenome molecular evolution on shorter time scales can be explained without invoking any adaptive benefit of mobile genes to their human hosts. However, a small subset of genes with distinct functions show dramatically different signature of molecular evolution, suggesting that selection acts at the level of gene function. Our results suggest that while host-related selective pressures may be strong over long evolutionary time scales, selection at the level of individual genes might predominate over shorter “human” time scales.

## RESULTS AND DISCUSSION

### Gene mobility correlates positively but not strongly with metagenomic coverage

To study pangenome evolution on time scales on the order of a human lifespan, we used an existing collection of mobile genes identified in 387 isolate genomes from the Human Microbiome Project (HMP) and 180 single-cell genomes from the Fiji Community Microbiome Project (FijiCOMP). Selected single-cell genomes came from 31 different genera and had less than 10% putative contamination called by CheckM^1,16^. The mobile genes were identified in genomic regions containing at least 500bp with >99% nucleotide identity over >50% of their sequence length between distantly related single-cell bacterial genomes (<97% identity in 16S rRNA), suggesting that HGT occurred within an individual human gut microbiome^1^. Ribosomal genes, which tend to be highly conserved, were excluded from this set of mobile genes as they could represent false-positive HGT events^1^. This procedure is strict, yielding likely true positive HGT events, at the expense of many false negatives^1,17^. We considered only genes with at least 10X metagenomic sequence coverage, and only metagenomes with at least 500 genes passing this coverage threshold. These filters yielded a total of 7,990 mobile genes out of the 37,853 genes present in the original dataset, and 175 out of 176 metagenomes, each from a different person from Fiji. We operationally defined gene mobility as the number of single-cell genomes in which a mobile gene was found. Gene mobility ranged from 1-16 species (mean = 2.73, standard deviation = 2.42; **Figure S1**) and is probably an underestimate of the true HGT rate because it was estimated from a limited sample (180 genomes) of the diversity in Fijian islanders’ gut. This could also be explained by small or incomplete assemblies of the single-cell genomes. Nonetheless, this dataset provides allows us to assess the balance of evolutionary forces in the pangenome on short timescales.

We began by asking whether our mobility metric behaves as expected in quantifying the spread of mobile genes in the gut. Assuming that genes with higher mobility will occur in more species, we expect them to be more deeply covered by metagenomic sequence reads. Consistent with this expectation, we found that a gene’s mobility is positively correlated with its depth of metagenomic read coverage (**Figure 1 and Table S1C**). The expectation of a positive correlation is not guaranteed because some mobile genes, such as selfish elements, have deleterious effects^18^ and can be subject to negative frequency-dependent selection ^12,19,20^ such that they are carried only by a fraction of individuals within a species, even if prevalent across species. The correlation between gene mobility and coverage is significantly positive in 169 out of 175 gut metagenomes (Bonferroni-adjusted p-value < 2.2×10^−16^), but the adjusted *R*^*2*^ and slope values are relatively modest (**Figure 1, Figure S2**). Varying selective pressures across mobile genes (*e.g.* deleterious effects and negative frequency-dependent selection) might by responsible for reducing the scaling between gene mobility and coverage, but not enough to flatten the relationship completely. We conclude that gene mobility, even if estimated from a relatively small sample of 180 gut bacterial genomes, behaves approximately as expected: generally leading to higher gene copy numbers.

**Figure 1.**
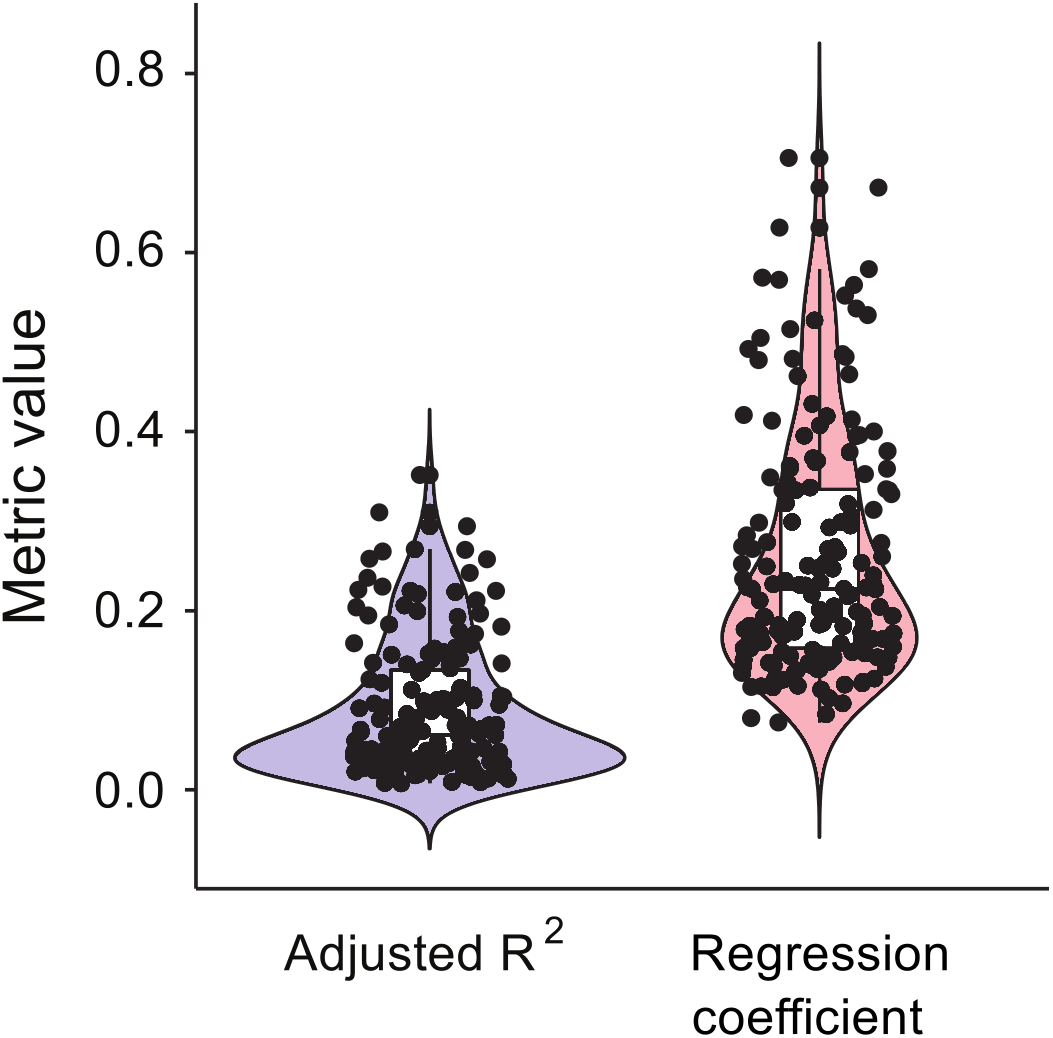
The correlation between gene mobility and metagenomic sequencing coverage is positive but widely variable. The boxplots and violin plots show the distributions of adjusted R^2^ values (blue) and slopes (red) across samples (individuals from Fiji) for the correlation between coverage (average depth per site) and gene mobility. The black dots represent the 169 samples (out of 175 tested) in which the correlation is significant (*t* test, Bonferroni-adjusted p-value < 0.05). Examples of this correlation in four randomly selected samples are shown in **Figure S2**.

### Estimating population genetic metrics from metagenomic data

The relationship between metagenomic coverage and gene mobility is generally positive but varies substantially across individuals (**Figure 1**). We therefore sought to ask whether this variation could be explained by either gene-specific factors (*e.g.* gene mobility and COG functional categories^1,19^) or human-specific factors, such age, diet, or social networks. Both gene-specific and human-specific factors are known to influence the patterns of mobile gene presence/absence across bacterial genomes^14^ and human hosts^1,20-22^, yet it is unclear if these patterns are explained by selection or drift. Here, we used the tools of population genetics to study molecular evolution of mobile genes based on their patterns of single nucleotide variants (SNVs) segregating in gut metagenomes. We quantified mobile gene sequence evolution using four population genetic metrics that detect selection and capture deviations from a neutral evolutionary model:

1. *θ*_π_, the nucleotide diversity calculated from the average number of pairwise nucleotide differences among metagenomic reads,
2. *θ*_*w*_, the nucleotide diversity calculated from the normalized number of segregating/polymorphic sites in metagenomic reads,
3. *Tajima’s D*, the normalized difference between *θ*_π_ and *θ*_*w*_, and
4. *dN/dS*, the ratio of nonsynonymous to synonymous substitution rates, measuring selective constraints at the protein level.

We note that our estimate of *dN/dS*, based on mapping metagenomic reads that could come from the same or different species, is a mixture of within-species polymorphism (often called *pN/pS*) and between-species divergence (*dN/dS*), but we refer to this hybrid metric as *dN/dS* for simplicity. We further note that *θ*_π_ and *θ*_*w*_ are two different estimators of the population mutation rate, *θ* = 2*N*_*e*_µ, where µ is the mutation rate and *N*_*e*_ is the effective population size. This difference in the two estimators is captured by *Tajima’s D*. In particular, *Tajima’s D<0* indicates more low-frequency mutations than expected under a standard neutral model with no selection and a constant population size^21^. This genetic signature can be the result of a population expansion, purifying selection, or a very recent selective sweep. Conversely, *Tajima’s D>0* indicates more intermediate- or high-frequency mutations than expected under a neutral model (**Figure S3**). It can be explained by population contraction, balancing selection, or negative frequency-dependent selection.

The above metrics were calculated for every gene in each sample by mapping metagenomic reads and calling SNVs after applying a 10X sequencing coverage filter (Methods). Consistent with previous estimates across multiple kingdoms of life^22^, we observe that *θ*_π_ and *θ*_*w*_ distributions across samples span 3 to 4 orders of magnitude (**Figure S4**). Also consistent with previous estimates in bacteria over different time scales^3,7,23^, *dN/dS* tends to be less than one, suggesting the predominance of purifying selection at the protein level (**Figure S4**). Our estimates of these population genetic metrics from metagenomic data are thus within an expected range and appear to behave as expected.

### Population genetic metrics vary more across mobile genes than across host attributes

With these metrics in hand, we asked whether mobile gene evolution is mainly driven by bacterial- or human host-specific selective pressures. To do so, we determined whether population genetic metrics varied more across gene families or across individuals. We first compared distributions of pairwise differences for each metric using the Kolmogorov-Smirnov test, and found much greater variation between genes than between individuals (**Figures 2 and S4**). This result indicates that, on short time scales, the selective pressures quantified by the four metrics may be less affected by person-specific factors, such as lifestyle or social networks, than by gene functions within a microbial cell. In other words, although some mobile genes may enable adaptations to personalized factors such as diet^1^, sequence evolution is relatively unaffected by these factors on short time scales (within an individual). In contrast, population genetic metrics vary substantially more across genes, suggesting that selective pressures act predominantly at the level of gene function.

**Figure 2.**
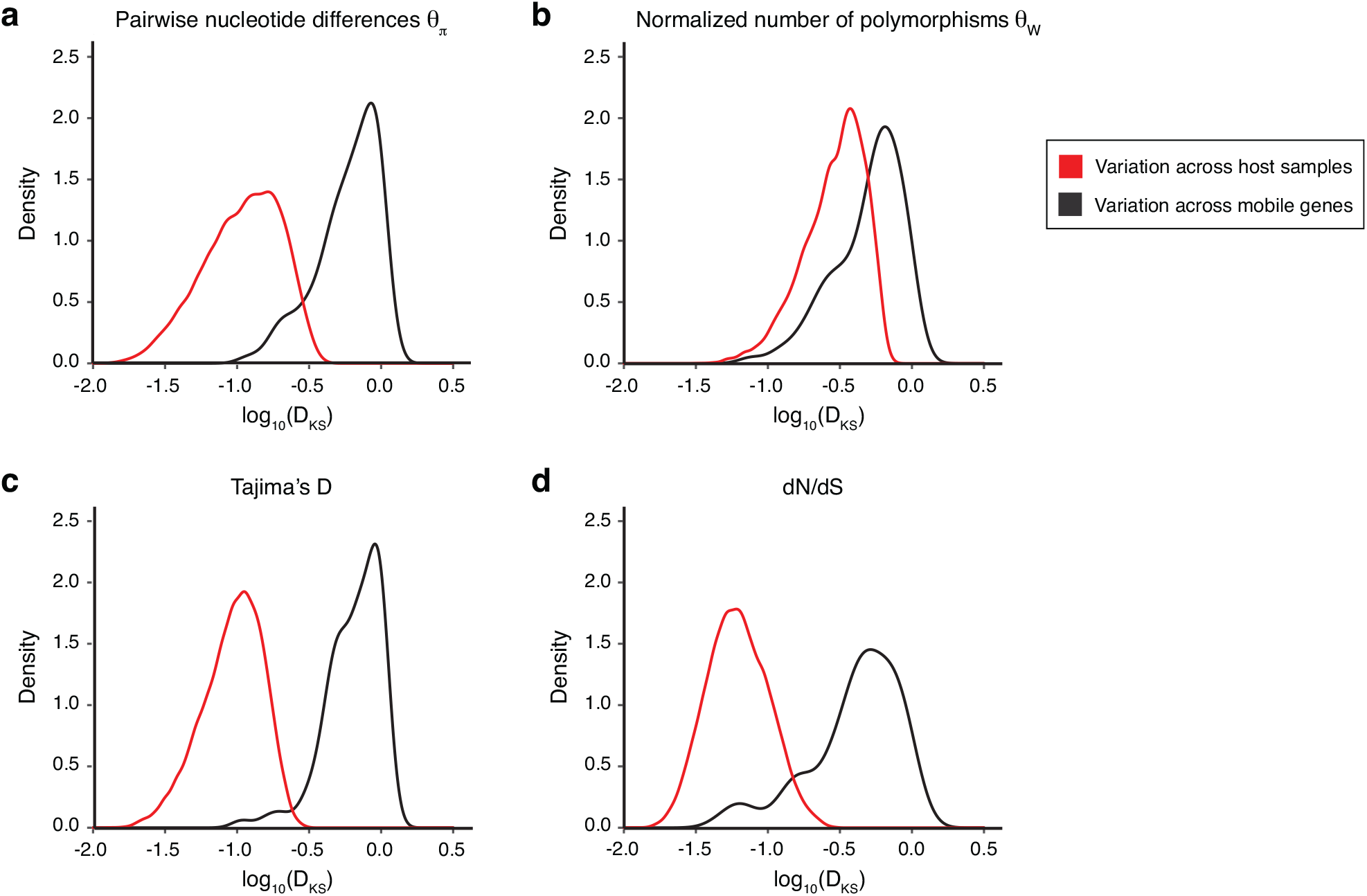
Mobile gene evolution varies more widely across genes than across samples (people). Each panel shows the distribution of the variation of population genetic metrics among samples (red) or among gene families (black) through the distribution of log_10_(*D*_*KS*_) statistics. The *D*_*KS*_ statistic from the Kolmogorov-Smirnov test measures the maximal distance between a pair of cumulative distributions – in this case, across either samples or genes. Panels a, b, c and d represent the variation of *θ*_π_, *θ*_*w*_, *Tajima’s D* and *dN/dS* respectively. We down-sampled the 37,853 genes to the same size as the number of samples set to avoid the potential bias toward more variation in the larger dataset of genes (999 sub-samples). This figure presents the result for 999 sub-samples of 175 genes and shows that there is more variation across genes than across samples/individuals for all the population genetics metrics (KS test, *P* < 2.2 x 10^−16^). See **Figure S4** for example distributions across genes and samples.

To validate that person-specific factors have weak effects on mobile gene sequence evolution, we used a linear regression where the continuous response variable is one of the population genetics metrics and the qualitative/categorical explanatory variable is a host attribute (Methods). Because the statistical significance of such an analysis is affected by sample size, we selected mobile genes with less than 30% missing values across the 172 samples for which metadata were available, for a total of 1333 tested genes. Host age and sex did not show any significant effects on mobile gene sequence evolution. However, a person’s household or village significantly influenced the evolution of just a few mobile genes (1.13% to 4.25% of the 1333 tested genes; **Figure 3A**). In this small subset of significant genes, the correlations between population genetic metrics and household (adjusted *R*^*2*^ ∼0.6 to ∼0.68) were stronger than correlations with village (adjusted *R*^*2*^ < 0.3), and these results were robust to varying the quality filters applied to the data (**Figure S5**). The small set of genes significantly influenced by household and village could be representative of very specific family/village selective pressures such as diet. Annotations of these genes show that they are involved in a set of functions involved in carbohydrates, lipids, secondary metabolites and ions transport or metabolism, and potential antibiotic resistance through ABC-type multidrug transporter system (**Tables S2**). Some of these functions are similar to those identified by Brito *et al.* as differentially abundant among villages or households^1^. Therefore, although village- or household-specific selective pressures do not explain much of the variation in population genetic metrics across genes, we cannot exclude rare instances in which social networks or lifestyles drive the evolution of few mobile genes over short time scales.

**Figure 3.**
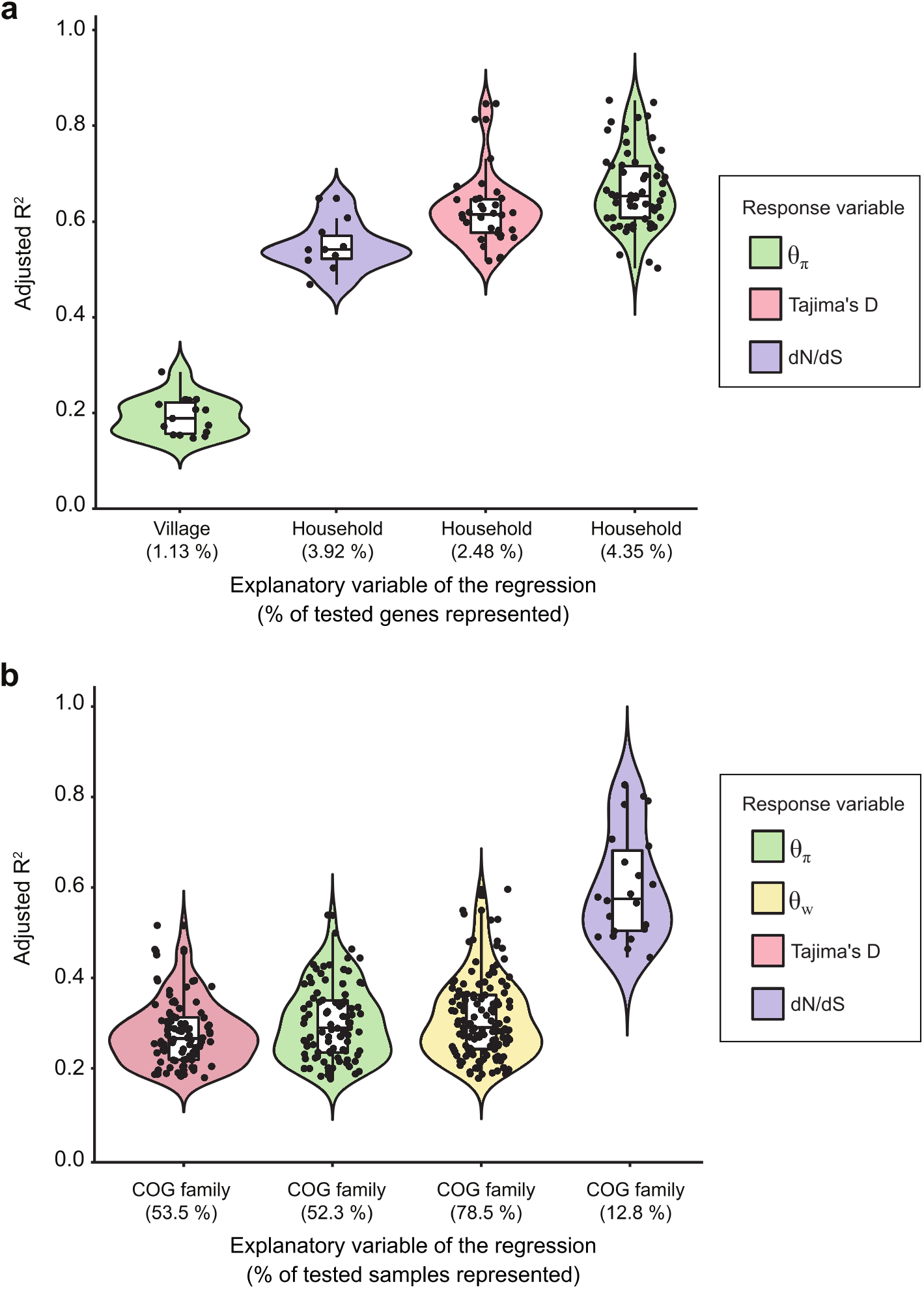
Gene function explains more variation in mobile gene sequence evolution than host attributes. A) Adjusted R^2^ values for the categorical regressions between population genetic metrics (color-coded) and host attributes. We only considered genes with at least 10X coverage in a sample, and we also required that mobile gene should have less than 30% missing values across samples, for a total of 1333 genes included in this analysis. The four strongest and most prevalent correlations between population genetics metrics and host factors are shown. Not shown are village significantly correlated with *θ*_*w*_ (0.15% of genes), *Tajima’s D* (0.15%) and *dN/dS* (0%) and household significantly correlated with *θ*_*w*_ (0.23%). Host age and sex did not show any significant effects on mobile gene sequence evolution. Each black point represents a mobile gene for which the categorical regression is significant. The percentage of significant genes out of the total number of genes tested is indicated in parentheses along the x-axis. For *dN/dS*, the sample size was reduced to n = 255 genes because an additional filter requiring mutations to be seen in a least 5 metagenomic reads was applied before computing *dN/dS*, which can other be sensitive to sequencing errors (Methods)**. B)** Adjusted R^2^ values of the categorical regressions between a population genetic metric and the COG family of the gene. Each black point represents a sample for which the categorical regression is significant. The percentage of significant samples out of the total number of samples tested is indicated in parenthesis along the x-axis. Only 172 out of 175 samples for which metadata was available are included in this analysis. We only considered genes with at least 10X coverage in a sample. We only included genes with a COG family annotation and required that each COG family be represented by at least 2 genes. Finally, we only included genes present in 30% or more of the samples, for a total of 512 genes included in the analysis.

Although host factors seem to have relatively little effect on the sequence evolution of most mobile genes on short time scales, selective pressures at the level of the genes might be more important. Indeed, we observed higher variations of population genetics metrics between genes than between samples (**Figures 2**), which could be explained by gene attributes such as their cellular function. To test this hypothesis, we used linear regressions between population genetics metrics and gene families based on the following set of conditions:

1. the gene should have at least 10X coverage to limit the impact of sequencing errors and to have confidence in the variant calling,
2. the gene should have an available COG family annotation (the explanatory variable in the regression),
3. the COG family should be represented by at least 2 genes within the dataset to avoid low sample sizes, and,
4. the mobile gene should have less than 30% missing values across samples, for a total of 512 tested genes.

In contrast to human factors, gene functions defined by COG families explained more of the variation in mobile gene sequence evolution across samples. For *θ*_*w*_, *θ*_*π*_, and *Tajima’s D*, COG families explained from ∼20% to ∼60% of the variance in >50% of the samples (**Figure 3B**). For *dN/dS*, COG families explained up to 83% of the variance in 12.8% of samples. To ensure that this result was robust to differential sampling of genes (n=512) and individuals (n=172) in this analysis, we downsampled to n=172 genes and confirmed that human host factors explain much less variation in mobile gene evolution compared to gene functions (**Figure S6**). A caveat of this analysis is that the strong explanatory power of gene functions on the population genetic metrics is based on COG functions being sufficiently well-represented in the dataset. Indeed, the strength of the correlations decreases with the stringency of the filters due to a decrease of sample size (**Figure S7**). As rarer genes (those present in fewer samples) are included in the analysis, fewer samples show significant correlations between *Tajima’s D* and COG function, going from ∼50% of samples when a gene can be absent in at most 30% of samples (**Figure 3B)** to ∼10% significant when a gene can be absent in 70% of samples (**Figure S7)**. It is therefore difficult to make conclusions about rare genes that are under-sampled in the dataset. However, for the more prevalent mobile genes, COG functions appear to explain much of their short-term molecular evolution.

### Higher gene mobility is associated with low-frequency SNVs in the gut microbiome

In addition to gene- or environment-specific selective pressures, the rate of HGT is also expected to affect mobile gene molecular evolution, as it allows genes to spread across different species, possibly altering their population size and thus the efficacy of selection^4,13^. To first order, each human host represents a distinct short-term evolutionary trial. Thus, to study the influence of HGT rate on molecular evolution within each of the human guts sampled, we correlated gene mobility with the population genetic metrics described above: *dN/dS, θ*_*π*_, *θ*_*w*_, and *Tajima’s D.* All correlation results reported below are robust whether or not we include gene length and coverage as covariates in linear regressions (**Figure S8**).

Using this regression approach, we first observed that the correlation between *dN/dS* and gene mobility was significant and positive in 144 out of 175 samples (**Figure S8**), but with a low average adjusted *R*^*2*^ of 0.03 (s.d. = 0.02). Although this result is consistent with slightly increasing positive or relaxed purifying selection with increasing gene mobility, we refrain from drawing strong conclusions due to the weak *R*^*2*^ values. We next observed that 159 out of 175 samples had a somewhat stronger significant correlation between *θ*_*w*_ and gene mobility (linear regression with Bonferroni-adjusted *p*-value < 0.05), and all the significant correlations were positive (mean adjusted *R*^*2*^ = 0.06; s.d. = 0.05). This is consistent with a model in which mobile genes accumulate SNVs that remain at low frequency (as measured by *θ*_*w*_, which is sensitive to these low-frequency mutations) as they spread across species. We also observed that *θ*_*π*_, which is more sensitive to intermediate-frequency mutations, decreases with gene mobility (**Figure 4A**). Among samples in which *θ*_*π*_ versus gene mobility regression results were significant (164 out of 175 samples with Bonferroni-adjusted p-value < 0.05), ∼95% of them exhibited this negative correlation (mean adjusted *R*^*2*^ = 0.08; sd = 0.05). As a result, *Tajima’s D* (which reflects the difference between *θ*_*π*_ and *θ*_*w*_) is significantly negatively correlated with gene mobility in ∼83% of samples (**Figure 4A**). Even if the *R*^*2*^ value are modest, we note that the trends are highly repeatable across samples. Reasons for the relatively low *R*^*2*^ values could include noise in the gene mobility metric (based on a small sample of genomes) and/or variable selective pressures across genes. There are several reasons for this enrichment of low-frequency SNVs (resulting in lower *Tajima’s D* values) in more mobile genes, including purifying selection keeping deleterious mutations at low frequency, recovery of new polymorphism after a recent selective sweep, or population expansion. This result suggests that HGT can spread genes across species faster than SNVs are able to rise to high frequency.

**Figure 4.**
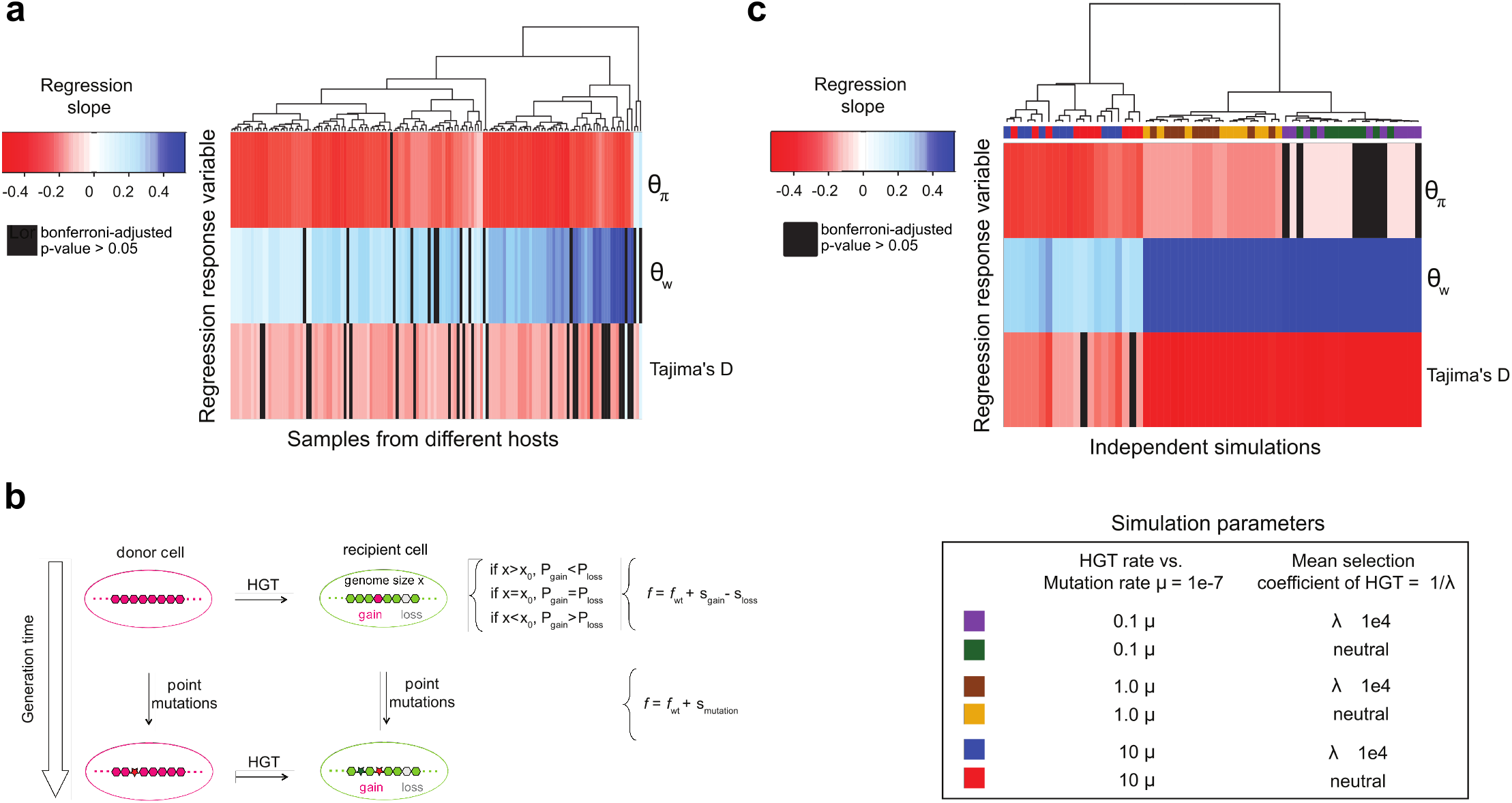
Gene mobility is negatively correlated with Tajima’s *D* in real and simulated microbiomes. A) Real data from Fiji. The heatmap shows the slope of a regression model in which either *θ*_*π*_, *θ*_*w*_ or *Tajima’s D* is the response variable and gene mobility is the explanatory variable (across samples). Regression p-values were obtained through a *t*-test. The heatmap contains non-significant regressions results after Bonferroni p-value filter (black), negative significant correlations (red) and positive significant correlations (blue). Data standardization was performed before each regression to respect the *t*-test’s assumption of normality. Heatmap rows and columns were clustered with Euclidean distance and complete linkage clustering. B) Representation of simulation events over two generations. In the first generation, a gene gain event occurs through HGT. Gene gain is represented by the transfer of gene from a donor cell to a recipient cell and increases the genome size of this recipient cell. The probability of future gene gain or gene loss events (*P*_*gain*_ and *P*_*loss*_ respectively) is determined by the difference between the current genome size of the cell (*x*) and the equilibrium genome size (*x*_*0*_). At equilibrium, the probability of gene gain and gene loss is the same by definition (*P*_*gain*_*=P*_*loss*_). An increase of genome size until it exceeds the equilibrium point (*x > x*_*0*_) leads to gene loss being more likely than gene gain (*P*_*gain*_*<P*_*loss*_). Gene gain also increases the fitness (*f > f*_*WT*_) of the recipient cell based on the selection coefficient of the transferred gene (*s*_*gain*_). In the model, each gene has its own selective coefficient which is drawn from an exponential distribution *exp(λ)* with an expected value of *1/λ*. Gene gain is either slightly beneficial or neutral in this model and has the opposite fitness effect of gene loss, which is slightly deleterious or neutral (-*s*_*gain*_ = *s*_*loss*_ where *s*_*gain*_ *>= 0*). Gene loss decreases the genome size of the target cell and in case this decrease leads to a smaller genome size than equilibrium, the probability of gene gain becomes higher than the probability of gene loss (*P*_*gain*_*>P*_*loss*_). Gene loss also decreases the fitness of the target cell (*f < f*_*WT*_) based on the selection coefficient of the lost gene (*s*_*loss*_). Finally, as represented in the second generation, mutations can also occur and change the fitness of the cell based on a selective coefficient (*s*_*mutation*_) which is drawn from a distribution (Methods). C) Simulated data. The heatmap shows the slope of a regression model in which either *θ*_*π*_, *θ*_*w*_ or *Tajima’s D* is the response variable and gene mobility is the explanatory variable (across simulation replicates). Simulations with different parameter for HGT rate and or distributions of selective coefficients (*s ∼ exp(λ)*) are color-coded (n=10 replicates per simulation).

### A subset of gene functions experiences a divergent regime of natural selection

Having established that *Tajima’s D* correlates negatively with gene mobility while coverage tends to correlate positively with mobility (**Figure 1**), we sought to determine if these general trends apply equally to all gene families. While the trends are significant across samples, the large variations observed across genes (**Figure 2**; **Figure S4**) could represent evolutionary regimes that are specific to some gene families. To test this hypothesis, we used linear mixed models with gene mobility as a predictor of either *Tajima’s D* or coverage as a response variable, while controlling for random variations across gut microbiome samples and allowing the response to vary across COG categories (Methods). This analysis was performed on genes with at least 10X coverage and available COG annotations (n= 3608 mobile genes).

As expected, based on the overall positive relationship observed (**Figure 1**), coverage and gene mobility are positively and significantly correlated across most COG categories (**Figure 5A**). COG category X (mobilome, prophages, and transposons) stood out as the strongest contributor to this positive relationship, consistent with this signal being driven by genes with the highest mobility. Removing sample identity or COG category from the linear mixed models significantly decreased the fit of the models, suggesting that they both significantly contribute to explaining variation in the mobility-coverage and Tajima’s D-coverage relationships (**Tables S1A and S1B**). We also confirmed that *Tajima’s D* is negatively correlated with gene mobility (**Figure 5A**), as observed in the regression analysis (**Figure 4A**). Deviations from this correlation could thus reveal signatures of selection that are specific to certain gene families. These COG categories for which *Tajima’s D* significantly increases with mobility, include P (Inorganic ion transport and metabolism), I (Lipid transport and metabolism), Q (Secondary metabolites biosynthesis, transport and catabolism), V (Defense mechanisms) and O (Posttranslational modification, protein turnover, chaperones), representing ∼30% of gene families (**Figure S9**). There are several explanations for why these gene families maintain or accumulate intermediate-frequency SNVs (*i.e.* an increase in *Tajima’s D*) while being transferred to many new species (**Figure 5B)**. The first explanation is a population contraction, or in this context, a reduction of the number of gene copies across species. However, this is unlikely for these subsets of genes because their coverage, which is a proxy of the relative abundance, increases with mobility. The second explanation is that these genes could be subject to species-specific selective pressures that fix mutations in some species but not others, resulting in intermediate SNV frequencies in the bulk metagenome. The third potential explanation is that negative frequency-dependent selection, which is thought to be an important force shaping pangenome evolution^20,24^, is acting on these genes, within species, between species, or both. Thus, the last two scenarios, which rely on the presence of distinct selective pressures on these subsets of genes, most likely explain how some mobile genes can maintain or accumulate intermediate-frequency SNVs as they spread across species.

**Figure 5.**
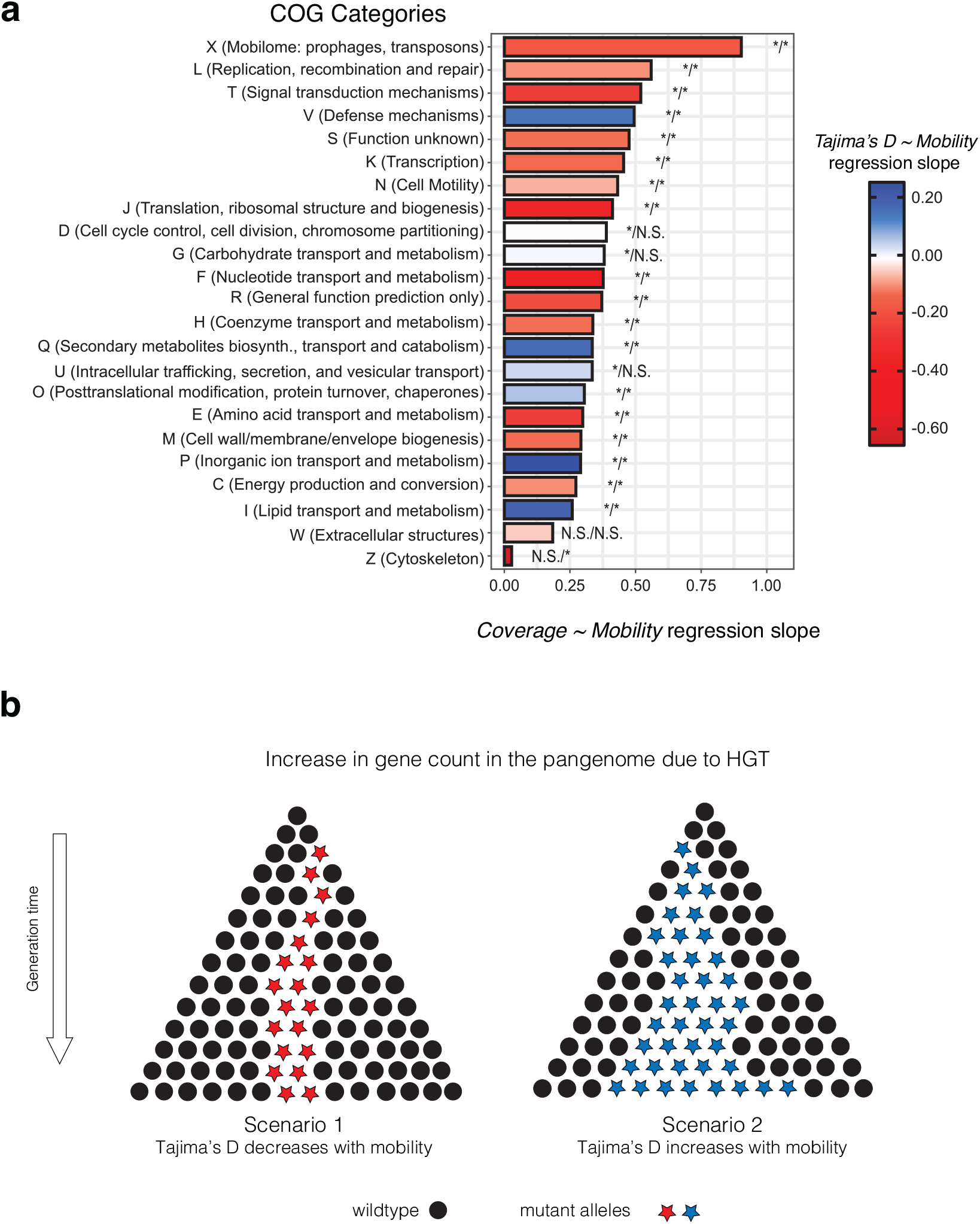
Gene mobility regressions reveal a minority of genes with distinct signals of selection. A) Linear mixed model regression slopes per COG category. This figure illustrate COG categories regression slopes for the linear mixed models “Coverage ∼ Gene mobility + Sample + COG category” and “Tajima’s D ∼ Gene mobility + Sample + COG category “with “Sample” and “COG category” being considered as random effects. Data were normalized using the Box-Cox transformation to ensure the condition of residual normality was accounted for before building the linear mixed model (Coverage Box-Cox λ = −0.01; Gene mobility Box-Cox λ = −0.005). We only used the 99.6% of Tajima’s D values that were negative and thus inversed their sign before applying Box-Cox transformation, which only works with positive values. We then performed the linear mixed model regression “−Tajima’s D ∼ Gene mobility + Sample + COG category “and inversed the sign of its slope (-*Tajima’s D* Box-Cox λ = 2). The sign of the slopes was consistent with simple linear regressions. The asterisks at the tip of each bar indicate the significance of the simple linear regressions “Coverage ∼ Gene mobility” and “Tajima’s D ∼ Gene mobility” respectively for the associated COG category (*=Significant; N.S. = Not Significant; Cut-off: Bonferroni-adjusted p-value < 0.05). B) Schematic of the evolutionary scenarios compared using linear regressions. Scenario 1 represents the situation in which mobile genes *Tajima’s D* is negatively correlated with gene mobility because HGT is faster than fixation of mutated alleles (red stars). Scenario 2 represents the situation in which *Tajima’s D* correlates positively with mobility. These genes maintain intermediate frequency mutations (blue stars) despite being frequently transferred to new species due to negative frequency-dependent selection or species-specific selective pressures that fix mutations in some species but not others. Note that the gene copies (dots or stars) illustrated here could come from members of the same or different species in the microbiome.

### Simple evolutionary simulations recapitulate the observed effects of HGT on mobile gene sequence evolution

To better understand potential mechanisms underlying the relationship between gene mobility and sequence evolution observed in the Fiji microbiome data, we implemented the explicit simulation of HGT and sequence evolution in SodaPop, a forward evolutionary simulation toolkit ^25^ (https://github.com/arnaud00013/SodaPop). Similar to Sela *et al.*, gene gain and loss are constrained to maintain genome size equilibrium and to have opposite fitness effects (**Figure 4B**)^7^. We used an updated version of the Sodapop model, which originally simulates protein sequence evolution with the distribution of fitness effects mutations derived from biophysics-based protein fitness landscapes ^25^. Briefly, we simulated a Wright-Fisher process for asexual populations ^25^ with 10 bacterial species. Each simulation included 5,000 cells in total, divided into 10 species, run for 10^5^ generations. Each gene has an explicit sequence which evolves by a Jukes-Cantor point mutation model ^28^, including synonymous sites that do not affect fitness and nonsynonymous sites with a distribution of fitness effects of which 30% are lethal^26^ (Methods). Genomes also experience HGT events, with explicit gene gain and loss events. The rates of these two events are updated at each generation for each cell to maintain an equilibrium around the genome size *x*_*0*_, set to 500 genes (**Figure 4B**) ^7^. Genomes larger than *x*_*0*_ are prone to gene loss, but genomes smaller than *x*_*0*_ are prone to gene gain. We also modeled gene gain and loss selection coefficients, specific to each gene and drawn from an exponential distribution with parameter *λ* (Methods). We kept simulated population sizes small due to memory limitations. To make sure this limitation does not cause excessive effects of drift (*e.g.* the accumulation of deleterious mutations leading to extinction, also known as Muller’s Ratchet ^27^) we forced species relative abundances to remain constant. We also set a relatively high mutation rate of 10^−7^ mutations per site per generation to compensate for the small population sizes and to ensure that enough mutations were generated in a reasonable number of generations. Genome size equilibrium was reached for every simulation, indicating robustness to variable starting conditions (**Figures S12**). Altogether, this model allows us to test if the relationships between gene mobility and population genetic metrics observed in the real data can be observed under varying rates of HGT and adaptive benefit of acquired genes.

We found that the simulation could recapitulate the major features observed in the real Fiji microbiome data without requiring that mobile genes provide adaptive value to a human host or to its bacterial genome. First, the simulations can recapitulate the shape of the observed distribution of gene mobility (**Figure S1**). A caveat is that simulations are far from including all the complexity of the gut microbiome, *i.e.* the number of species, population structures and other features not simulated, and the distributions were only compared for one illustrative set of input parameters (**Figure S1**). Thus, we do not claim that our model can provide a precise quantitative description of gene mobility in the gut microbiome, but rather that it can recapitulate the major qualitative features.

Second, the simulations recover the positive correlation between gene mobility and census population size (metagenomic coverage) observed in the real data (**Figure 1**). The positive correlation was always stronger in the simulations (mean adjusted *R*^*2*^ of 0.705 across all parameter settings, standard deviation = 0.190) compared to the real data (mean adjusted *R*^*2*^ of 0.085 across all parameter settings, standard deviation = 0.076). This suggests that factors not included in the model, such as negative frequency-dependent selection and noise in the gene mobility metric, reduced the strength of the correlation in the real data. The positive correlation was stronger in simulations with relatively lower HGT rate but was largely unaffected by whether HGT events were neutral or adaptive to host cell fitness (**Table S3**). This suggests that relatively high HGT rates could also explain the weaker correlation between gene mobility and coverage observed in the real data.

Third, we assessed whether the simulations could reproduce the observed correlations between population genetics metrics and gene mobility. Simulations recapitulated most of the observed effects of HGT on nucleotide diversity in real data. Specifically, *Tajima’s D* correlates negatively with gene mobility in simulations, with a median adjusted *R*^*2*^ of 0.32 (mean = 0.23; sd = 0.13) compared to a median adjusted *R*^*2*^ of 0.01 (mean = 0.01; sd = 0.01) in the real data and reproducible across ∼87% of simulations compared to ∼83% of the samples in the real data (**Figure 4**). The variation in this correlation is explained more by HGT rate than by HGT fitness effects (neutral or adaptive selective coefficients on gene gain/loss). This can be seen in the heatmap, in which simulations cluster by HGT rate rather than HGT fitness effect (**Figure 4C**). Along the same lines, we performed a K-S test on the slopes of the regression between *Tajima’s* D and mobility and observed that this slope varies more because of HGT rate than HGT fitness effect (**Figure S10**). Simulations also predict that *dN/dS* also correlates positively but weakly with mobility, but only at intermediate HGT rates (**Figure S11)**. A similar pattern is observed in the real data, in which *dN/dS* correlates weakly with mobility (**Figure S8**). Overall, real microbiome data is recapitulated by our simple evolutionary model, which includes only selection for a stable genome size, without the need to invoke adaptive advantage of mobile genes to their bacterial genomes, or to include any human host factors whatsoever.

## CONCLUSION

Pangenome evolution has been studied primarily on long evolutionary time scales by comparing relatively distantly related genomes. Studies of these long time scales have largely concluded, although with some debate, that pangenomes are predominantly adaptive – that selection plays a bigger role in pangenome evolution than drift. Here we have refocused the study of pangenome evolution to shorter time scales, that is within individual gut microbiomes in which gene transfer events likely occurred within a human lifespan. Based on microbiome data from a Fiji cohort, we found that mobile gene sequence evolution is more influenced by selective pressures at the level of gene function than at human host level. Of course, there were many unmeasured human host factors which could impose selective pressures that we were unable to study. However, complementary evolutionary simulation results showed that mobile genes need not provide any special adaptive value to their human host or microbial genomes in order to recapitulate the qualitative patterns of molecular evolution observed in the real data.

These observed patterns of molecular evolution based on population genetic metrics provide clues about the balance of evolutionary forces acting on mobile genes in microbiomes within a human lifespan. We found that most genes accumulate low-frequency mutations as they spread within and between bacterial species. One interpretation of this result is that most mobile genes are under purifying selection to maintain a conserved function, even as they spread across species, such that most mutations are deleterious and kept at low frequency. Another non-exclusive interpretation is that low-frequency mutations could also represent rapid spread of a gene, before mutations are able to rise to higher frequency. In contrast, a minority of genes involved in few specific cellular functions, such as defense mechanisms (COG category V), accumulate intermediate frequency alleles as they spread in new species, possibly due to negative frequency-dependent selection within species and/or fixation of beneficial mutations within some species but not others. Further investigation will be needed to explore the nature of these variable selective pressures.

Similarly to Bobay and Ochman (2018), we observed a very weak correlation between gene mobility and *dN/dS*, which measures selection in protein-coding regions. Bobay and Ochman (2018) attributes this trend to a nearly neutral model of pangenome evolution, *i.e.* drift-barrier evolution. This assumption that most accessory genes are slightly beneficial could explain why a mixture of neutral and adaptive patterns are evident throughout our analysis. Further work is needed to test the validity of this model in additional datasets.

Thus, pangenome evolution is the product of a fine balance between drift and selection, which can shift depending on the time scale and level of biological organization, from gene to genome to community. In the gut microbiome of a single person, the time scale of evolution may be too short to easily resolve the balance between drift and selection. Indeed, on very short time scales during which mutations could still be segregating and HGT occurs more frequently than mutation fixation, slightly adaptive genes that have been recently transferred could be largely influenced by drift because of their small *N*_*e*_, such that their adaptiveness could be effectively detected only on long time scales, while drift might decide their fate on shorter time scales. In this context, it is not surprising that simulations identified HGT rate, but not selective coefficients, as an important driver of molecular evolution. This model seems to fit some other bacterial genomic datasets^11,23^ but awaits formal testing. Finally, we suggest that future work on pangenome evolution should try to understand what factors control shifts in the drift-selection balance and its interplay with species ecology (*N*_*e*_, species lifestyle, etc.) and gene ecology (*i.e.* gene function, to what extent are genes selfish or cooperative within a genome, etc.), which is probably more informative than simply settling for either an adaptive model or a non-adaptive model.

## METHODS

### Population genetics of Fijian gut microbiome mobile genes

The Fiji Community Microbiome project provides open access to metagenomes from the gut microbiomes of 176 individuals. For each of these individuals, we mapped metagenomic sequence reads to a set of 37,853 mobile genes previously defined as follows from bacterial whole genome sequences from the Human Microbiome Project (HMP) and FijiCOMP. To be considered mobile, pairs of genes 500bp or longer had to share >99% nucleotide identity between isolate or single-cell genomes with <97% identity in the 16S rRNA gene^1^. This procedure selects nearly identical genes present in distinct species or genera as candidates for very recent HGT, likely within an individual gut microbiome^1,17^. An additional filter was applied to remove potential false-positive HGT events from highly conserved ribosomal proteins, and to keep only reads that aligned with 99% identity across >= 50% of their own length ^1^. From the mappings, we used Anvi’o to report Single Nucleotide Variants (SNVs) (--min-coverage-for-variability 10 --min-contig-length 50), followed by a pipeline to compute population genetics metrics (*θ*_*π*_, *θ*_*w*_, *dN/dS* and *Tajima’s D*) based on the SNVs. The pipeline scripts are available at https://github.com/arnaud00013/Fiji_Mobile_Gene_Specific_PopGen_scripts. The Anvi’o SNV calling module^28^ has the advantage of being fast and simple to use, can be executed in parallel (High-Performance Computing), and has filters to control minimum gene coverage or mutation frequency. For each sample mapping, a gene was retained if its mean site depth was >= 10. Only one sample was excluded for having less than 500 genes passing the site depth filter, reducing the sample size to 175 metagenomes. Among all samples, 7990 unique genes were conserved after applying the site depth filter. Finally, mobile gene COG annotations, available in the FijiCOMP data (http://fijicomp.bme.cornell.edu//), were used to define two level of gene functions: COG gene family (which is more specific), and COG category (which is more general).

### Detecting selection by *dN/dS*

*dN/dS* is the non-synonymous to synonymous mutations per site ratio. Different methods have been developed to estimate *dN/dS* with the common purpose of inferring selection in protein-coding genes^29^. More precisely, *dN/dS* can detect purifying selection (*dN/dS*<1), neutral evolution (*dN/dS* ≈ 1) and positive selection (*dN/dS* > 1). Because we are working with metagenomic gene variants, we defined our own estimator of *dN/dS*:

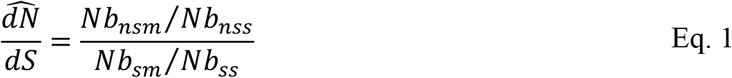

where Nb_nsm_ is the number of non-synonymous mutations (SNVs), Nb_nss_ is the number of non-synonymous sites, Nb_sm_ is the number of synonymous mutations (SNVs), and Nb_ss_ is the number of synonymous sites.

### Measuring mobile genes nucleotide diversity at metagenomic level

Because mobile genes are by definition present in multiple species, we calculated population genetic metrics based on all reads from a metagenome that map to a particular mobile gene. Based on these mapped reads, we calculated *Tajima’s D* ^21^, which measures the difference between average per-site pairwise nucleotide differences (*θ*_*π*_) and the normalized number of polymorphic sites (*θ*_*w*_) :

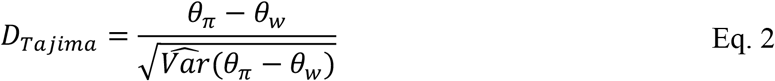

where the 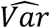 denotes the expected sampling variance of (*θ*_*π*_ − *θ*_*w*_). For each sample, we estimated mobile gene nucleotide diversity from sequence variants detected in the mapping between metagenomic reads and mobile gene reference sequence from FijiCOMP as follows:

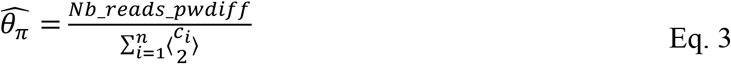

where n is the gene length, c_i_ is the depth of the site i of the gene and Nb_reads_pwdiff is the number of pairwise nucleotide differences, and

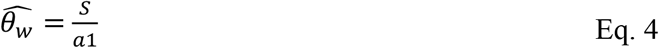

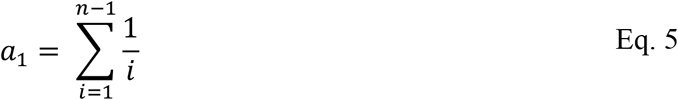

where a_1_ is a normalizing factor that represents the sample size (n). Usually, *Tajima’s D* is estimated from a multiple alignment between gene alleles. The sample size used to estimate the normalizing factor a1 is the number of alleles. Here we use the average depth of coverage at polymorphic sites as an estimator of the sample size n.

### Effect of gene mobility on metagenomic coverage

We operationally defined gene mobility as the number of single-cell genomes in which a mobile gene was found and tested if this metric behaves as expected in explaining gene frequencies in metagenomes. More precisely, we correlated gene mobility with metagenomic coverage with the expectation that more mobile genes occur in multiple species and should thus be more deeply covered by metagenomic sequence reads. Linear regression analyses and t-tests were calculated using the R function “summary.lm()” ^30^. Data standardization was performed before each regression to respect the t-test’s assumption of normality.

### Assessing variation in sequence evolution across genes and across individuals

To determine whether mobile gene evolution is driven more by gene-specific factors or by host attributes, we first compared the variation of mobile genes nucleotide diversity (and other population genetic metric described above) across genes vs. across samples through the Kolmogorov-Smirnov test (KS test). The KS test involves a statistic *D*, which measures the maximal distance between a pair of cumulative distributions. We downsampled the mobile genes to the same size as the number of samples to avoid the potential bias due to different sized datasets and repeated this for a total of 999 resamples. We performed this series of KS test with the function ks.test() from the R package “stats” ^30^.

### Gene function and human host (individual) attributes as predictors of mobile genes evolution

To determine whether mobile gene evolution is driven more by gene function or host attributes, we performed linear regressions between a continuous response variable and a qualitative/categorical explanatory variable, which we will refer as a factor. Regressions between a quantitative continuous variable, e.g. *Tajima’s D*, and a factor, e.g. gene function family, requires transforming the factor as it cannot be integrated into a regression equation in its original form ^30^. We therefore used the R contrast function “constr.sum()” to transform factors ^30^. This transformation allows the regression coefficients to represent how each level/state of the factor differ. Then, we assess the significance of the regression model with a non-parametric (permutational) ANOVA ^31^. This test makes random permutations of the response variable between the different groups/levels of the factor, and estimates the *p*-value as the proportion of permutations with an F-statistic greater than or equal to that observed in the real (unpermuted) data. This test was implemented in the R library “lmPerm” (v.2.1.0) ^30^.

For host attribute correlations with population genetic metrics, we focused on 172 samples with available metadata. Metadata about these samples were extracted from Brito et al. (2016) and NCBI accession numbers of the corresponding stool metagenomes are publicly available at http://fijicomp.bme.cornell.edu//data/FijiCOMPmetagenomicsamples.xlsx. Mobile genes selected for this analysis needed to respect the following conditions: (1) the gene should have at least 10X coverage to limit the impact of sequencing errors, and (2) mobile gene should have less than 30% missing values across samples, for a total of 1333 tested genes.

As for linear regressions between population genetics metrics and gene families, we selected genes based on the following set of conditions : (1) the gene should have at least 10X coverage to limit the impact of sequencing errors, (2) the gene should have available COG family annotation, (3) the gene COG family should be represented by at least 2 genes within the dataset and (4) the mobile gene should have less than 30% missing values across samples, for a total of 512 tested genes. The first two filters are the basic requirements for doing these regressions analyses. However, the 3^rd^ and 4^th^ filters were chosen respectively to avoid the effects of small sample size for COG families that are underrepresented in the dataset, and to handle missing values caused by gene absence across sample or genes with low coverage in gut metagenomes.

### Effect of HGT on sequence evolution

To determine the impact of HGT on mobile gene sequence evolution, multiple linear regressions were performed. In these multiple linear regressions, coverage, Gene Mobility – the number of species in which a mobile gene has been identified when looking for HGT events – and gene length were the explanatory variables and the various population genetic metrics were the response variables. We used the lm() function in R to remove collinearity with QR-decomposition/Gram-Schmidt orthogonalization. Thus, it is possible to assess the effect of Gene Mobility on each population genetics metrics while controlling for the effect of potential confounders like coverage and gene length. For each response variable Y tested (*θ*_*π*_, *θ*_*w*_, *dN/dS* and *Tajima’s D*), there are two regression models:

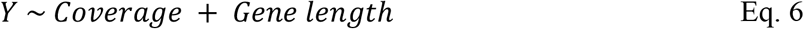

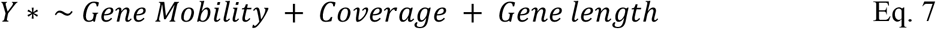

The asterisk represents the fact that the regression controls for the effects of coverage and gene length, which increase the chance of observing sequencing errors. The adjusted R^2^ of a correlation represents the proportion of variable Y variance that is explained by the regression model with a correction for the number of explanatory parameters included in the model (k) and the sample size (n):

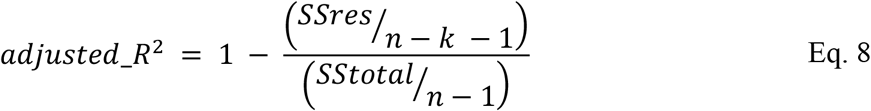

where SSres is the residual sum of squares and SStotal is the fitted data sum of squares. The type of correlation (positive or negative) can be determined by the regression coefficient. The reproducibility of the regressions was measured by the number of samples in which the correlation is significant.

### Variation across COG categories

To assess how the relationships between gene mobility and *Tajima’s D* or coverage varied across COG categories, we considered 22 COG categories ^32^. We then used linear mixed models, through the R package lme4, to study the effect of gene mobility on coverage and *Tajima’s D* across COG categories ^33^. A linear mixed model allows to build a linear model between the response variable and the fixed effects while controlling for random effects. In the regression model, fixed effects are explanatory variables for which we want to know the relationship with the response variable. Random effects are grouping factors that explain random variance of the relationship between the response variable and the fixed effects across a finite number of different groups. To control for random effects, the algorithm builds a linear model for each group. In the two regression models, “COG category “and “Sample” were included as random effects:

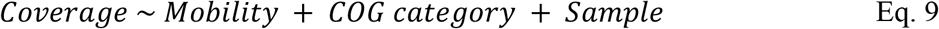

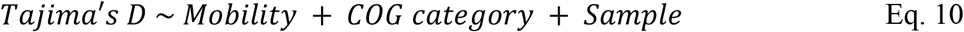

We can then test the significance of “COG category” for the regression model using a permutation ANOVA ^31^. The advantage of such test is that it is non-parametric, making no assumptions about the distribution underlying the data. For both regressions, we conducted 99,999 permutations of the response variable between COG categories and then calculated the F-statistic of the regression after each permutation. Next, we calculated the F-statistic of the original regression and calculated the *p*-value as the proportion of permuted data regressions that gave an F-statistic greater than or equal to the F-statistic from the real (non-permuted) data.

Additionally, using the R function anova(), we performed likelihood ratio tests between each linear mixed model and their nested models to test the significance of each random factor, i.e. “COG category “and “Sample “^30,34^. Each nested model was obtained by removing one random factor at a time, thus creating two nested models per response variable Y:

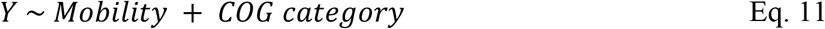

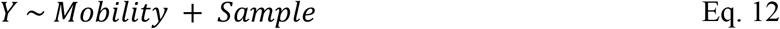

The likelihood ratio test compares the likelihood of a nested model to the likelihood of the full linear mixed model, with the assumption that the test statistic follows a Chi-square distribution. Thus, we can create each nested model by the removal of a single random factor from the full linear mixed model and assess the significance of both random factors using a *p*-value from the Chi-square distribution ^34^.

### Simulation of pangenome evolution

We simulated Sela, Wolf and Koonin’s prokaryotic genome size evolution model with few changes, using the SodaPop simulation tool ^7,25^. In this model, the selective advantage of gene gain, *i.e.* the advantage of having x+1 genes instead of x genes, depends of the genome size, which is measured by the number of genes in the genome (x). Selection coefficients for gene loss have the opposite sign as gene gain; thus, gene gain is slightly beneficial while gene loss is slightly deleterious^7^. The selection coefficient of gene gain and gene loss can thus be described by the following formula:

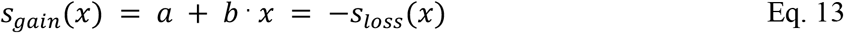

where *s*_*gain*_ is the selection coefficient of gene gain through HGT, “*a*” is a constant input parameter of the simulation that allows to improve the fit of the linear expression with the real data, “*b*” is a constant input parameter that represents the benefit or cost associated with the gain of a single gene, x represents genome size (number of genes), and *s*_*loss*_ is the selection coefficient of gene loss. We modified this formula to simulate a model where each gene has its own constant selective advantage regardless of genome size (x). To do so, we only needed to set the condition b = 0. This change allowed us to reproduce the shape of gene mobility distribution in simulation (**Figure S1**). In this case:

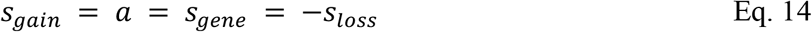

where *s*_*gene*_ ∼ Exp(λ), λ is an input parameter of the simulation, and 1/λ represents the expected value of the exponential distribution of selection coefficients.

In the model, genome size (x) influence gene gain rate and gene loss rate. Indeed, the more genome size increases, the more gene gain rate decreases, and the more gene loss rates increases to find an equilibrium around a certain genome size x0. Therefore, when genome size (x) is smaller than genome size at equilibrium (x0), the cell has a higher probability of gene gain than loss. To consider the stochastic component of evolution, the cells and genes that are involved in each gain or loss events are randomly selected. Also, the number of gain or loss events are drawn from a Poisson distribution with the gain and loss rates as follows:

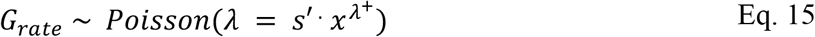

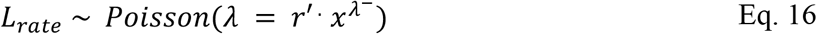

where *G*_*rate*_ is the gain rate, i.e. the number of gene gain events per generation, *L*_*rate*_ is the loss rate, i.e. the number of gene loss events per generation, and r’, s’, *λ*^+^ and *λ*^−^ are simulation input parameters that allow to tune the gain and loss rates.

We implemented this model in the SodaPop software, which simulates a Wright-Fischer process for asexual populations ^25^. In SodaPop, the mutation model is equivalent to Jukes-Cantor in which all single nucleotide occur at the same constant rate^35^. We also implemented a distribution of non-synonymous mutation fitness effect in which 30% of mutations are lethal, as previously reported in literature^26^, and 70% are drawn from a normal distribution, N(µ=-0.02, σ=0.01). Synonymous mutations are all considered neutral unless the user provides data on species codon usage and the related fitness effects. SodaPop also offers flexibility in the initial setup of the simulation^25^. We created scripts to facilitate the creation of the simulation starting conditions (https://github.com/arnaud00013/SodaPop/tree/Sodapop-pev/tools). The scripts allow to define each species abundance, gene content, and to define the genes that are mobile (https://github.com/arnaud00013/SodaPop/blob/Sodapop-pev/tools/Setup_SodaPop_with_PEV.py). Mobile genes can be transferred and lost while core genes and accessory genes (defined at the start of the simulation) can only be lost. For each set of simulations sharing the same input parameters, we ran 10 replicates. Each simulation included 5000 cells, 10 species, 500 genes per cells at equilibrium and a simulation time of 10^5^ generations and a timestep of 10^4^ generations to save simulation data. Population size is small in simulation because of hardware memory limitations. To avoid undesirable effects, like Muller’s Ratchet, we maintained species abundance constant. We also established a relatively high mutation rate on the order of 10^−7^ mutations per site per generation to compensate for small population sizes. Genome size equilibrium was reached for every simulation and the model is thus robust to the initial conditions (**Figures S12**). The software is available on GitHub (https://github.com/arnaud00013/SodaPop).

## ACKNOWLEDGEMENTS

We would like to thank Compute Canada for allocated resources, and Louis-Marie Bobay and Gavin Douglas for constructive comments. BJS was supported by an NSERC Discovery Grant. AWRS acknowledges funding from a Canada Research Chair Tier 2 and an NSERC Discovery Grant. AG is supported by an FRQNT scholarship.

