## Supplemental Figures and Tables for "Pangenome sequence evolution within human gut microbiomes is explained by gene-specific rather than host-specific selective pressures"

### SUPPLEMENTARY MATERIAL

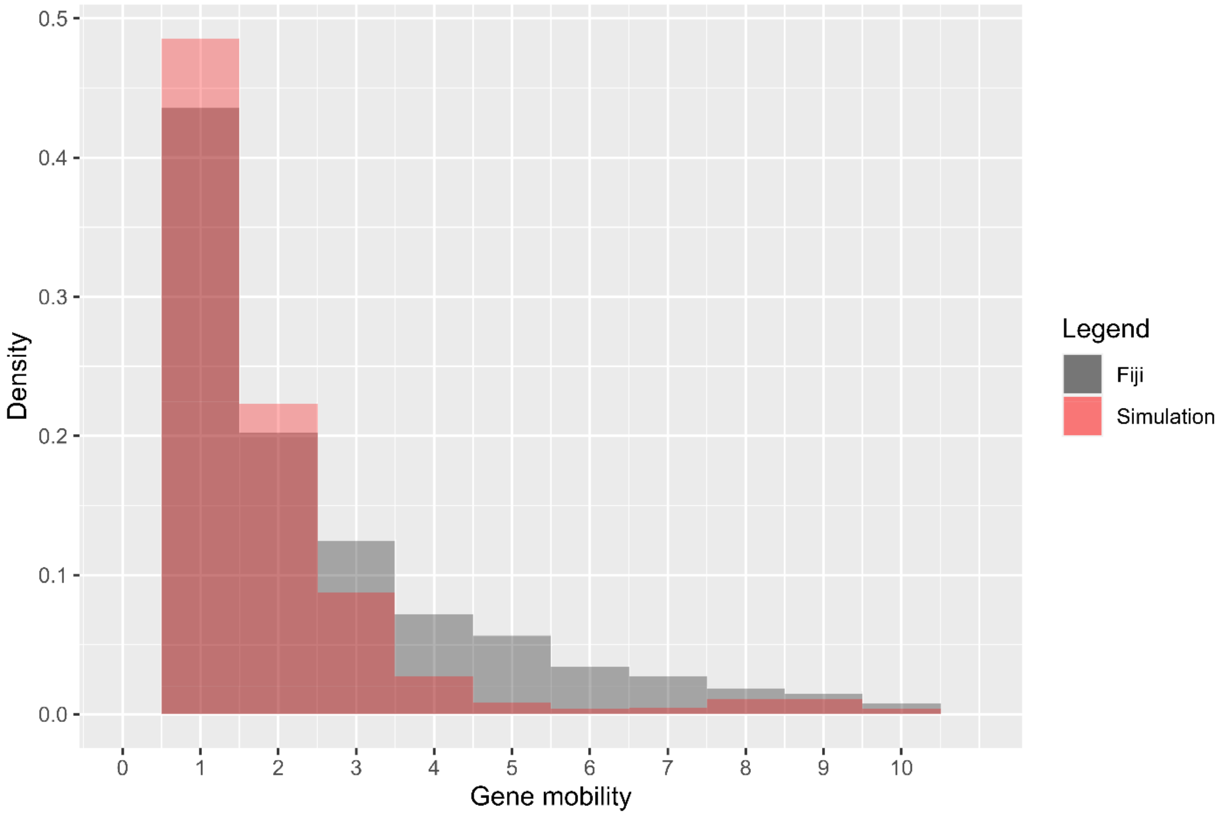


**Figure S1. Gene mobility distribution in simulation vs in Fiji dataset**.

Simulations in which HGT is slightly adaptive and the HGT selection coefficient is exponentially distributed produce a gene mobility distribution with a similar shape than the one observed in the Fiji dataset. However, their quantitative similarity is not significant (Kolmogorov-Smirnov test p-value $< 0.05$) and the range of mobility in simulation is [1,10] while it is [1,16] in the Fiji dataset so the distribution is truncated here to a maximum of 10 species. The simulation presented (red) included 5000 cells, 10 species, 500 genes per cells at equilibrium, a simulation time of $10^5$ generations, HGT rate = 10µ and HGT selection coefficient parameter $\lambda$ = 1E5 (HGT is slightly adaptive; *s* = 1E-5 in average).

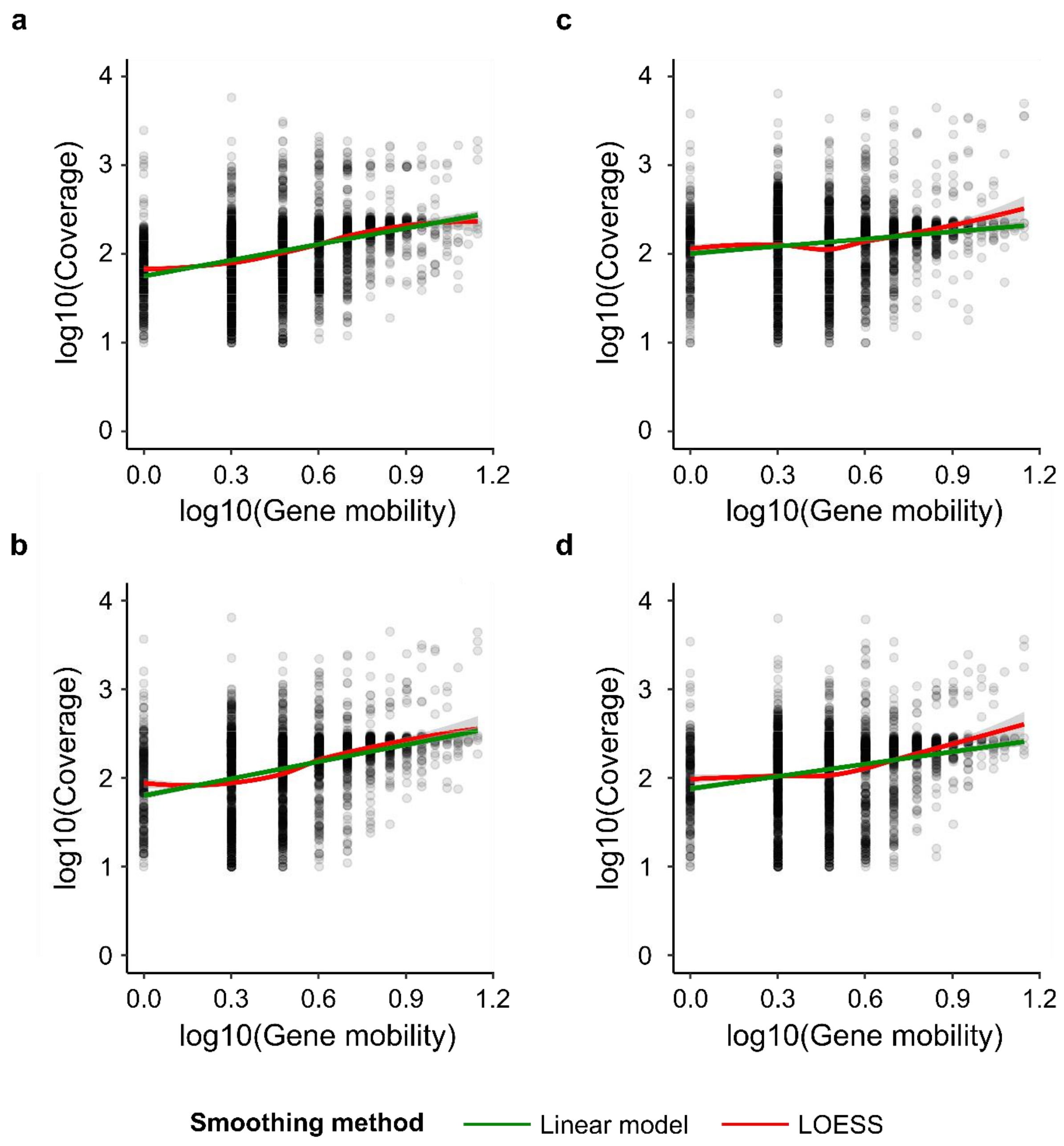


**Figure S2. Coverage in function of gene mobility in log10 scale across 4 samples**.

These figures show examples of the correlation between a mobile gene coverage and its mobility in log10 scale for samples a) G30512, b) G30771, c) G30520 and d) G30804. The correlation observed in these samples is positive, which is consistent with the main trends. Additionally, coverage and mobility range are shown in the figure. Coverage is always greater than 10x, which is a minimal requirement for variant calling. Finally, linear model (green) and LOESS (red) smoothing curves are represented by the colored lines.

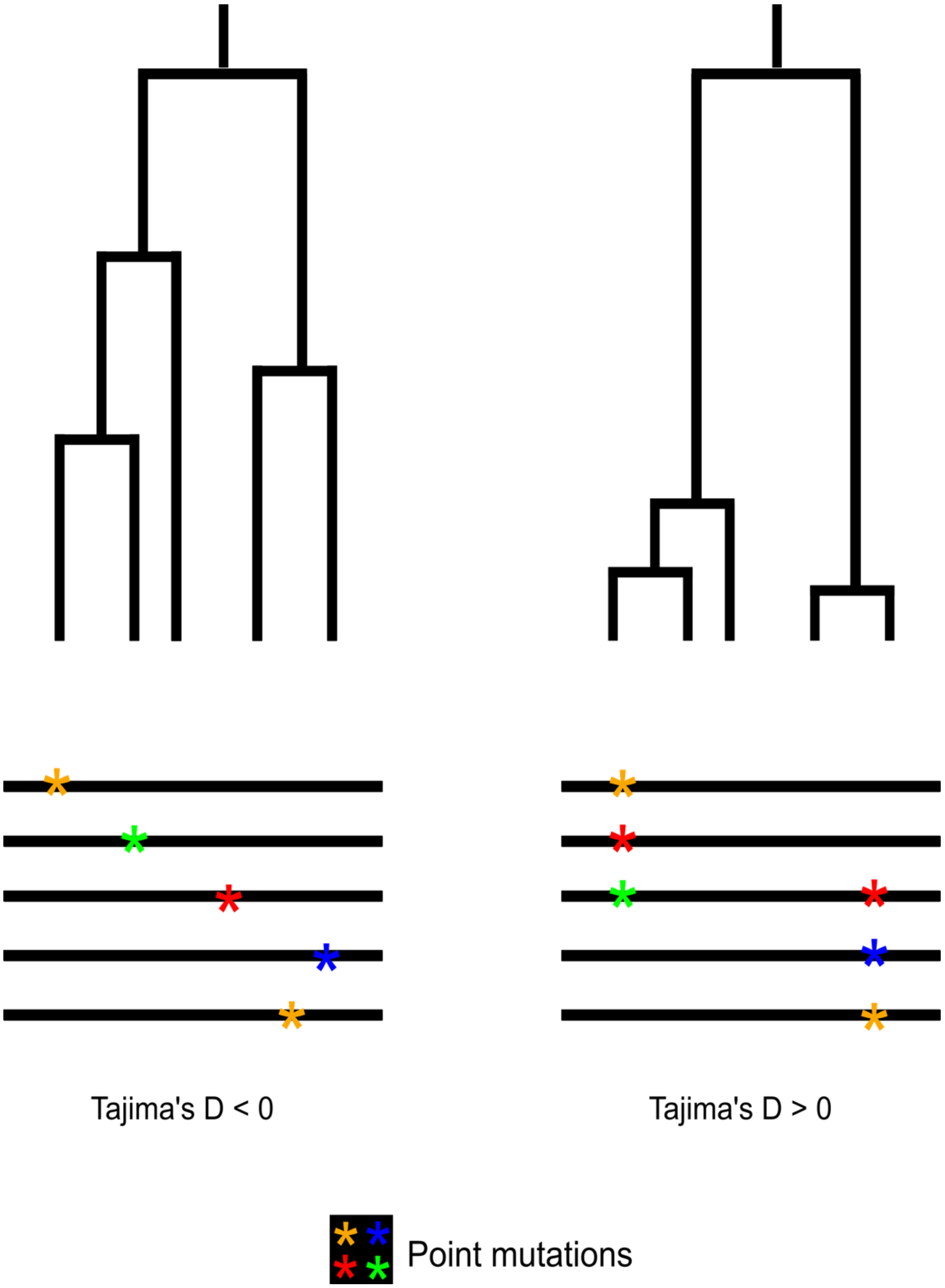


**Figure S3. Underlying logic of *Tajima's D*.** *Tajima's D* measures the difference between average per-site pairwise differences of gene alleles ($\theta_\pi$) and the normalized number of polymorphic sites ($\theta_w$). It takes negative values when there are more low-frequency mutations than expected under a standard neutral model with no selection and a constant population size [20]. In a phylogenetic tree of the gene alleles, this would result in high divergence and long branches as the low-frequency mutation are rarely shared across alleles and occurs at different polymorphic sites most of the time, which can be observed in the corresponding alignment. This genetic signature can be the result of a population expansion or a very recent selective sweep. Positive *Tajima's D* values are observed when there are more intermediate/high frequency mutations than expected under a neutral model [20]. In a phylogenetic tree of the gene alleles, this would result in low divergence and shorter branches as the intermediate/high frequency mutations are frequently shared across alleles and occurs at few polymorphic sites, which can be observed in the corresponding alignment. Positive values of *Tajima's* D can be explained by population contraction or negative frequency-dependent selection.

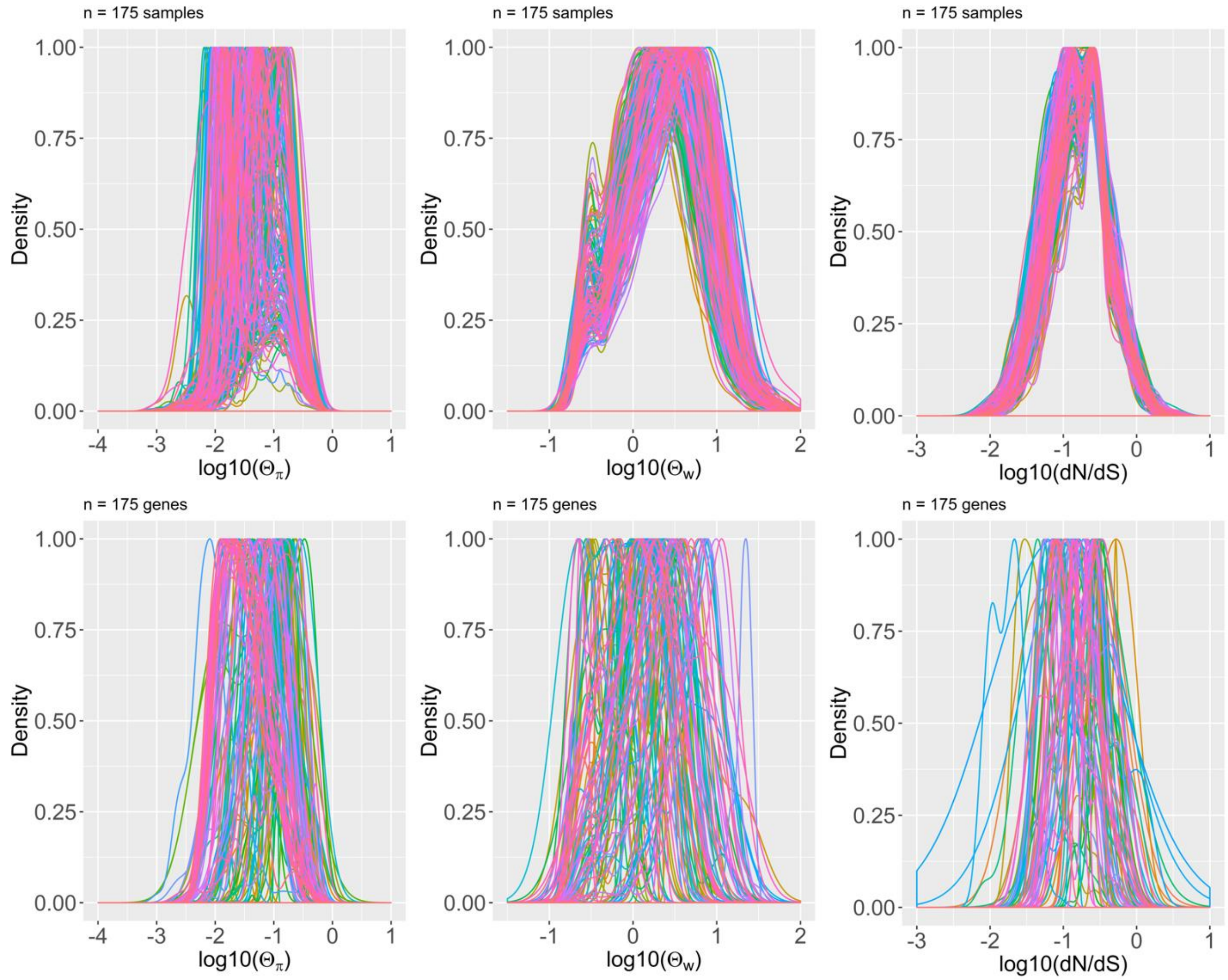


**Figure S4. Population genetics metrics distributions vary more across genes than across samples (people)**

We measured and compared the variation of mobile genes population genetics metrics across samples/individuals and across genes. The top row represents the distributions of *dN/dS*, $\theta_\pi$ and $\theta_w$ (logarithmic scale) across samples and the bottom rows represents the distribution of these descriptors across genes (downsampled to 175 to match the sample size). Each column represents one of three replicates random downsamplings. It can be visually seen that the distributions are more variable across genes (bottom row) than individuals (top row), and this is quantified by the Kolmogorov-Smirnov *D* statistic, computed for each of 999 replicate downsamplings to produce **Figures 2**.

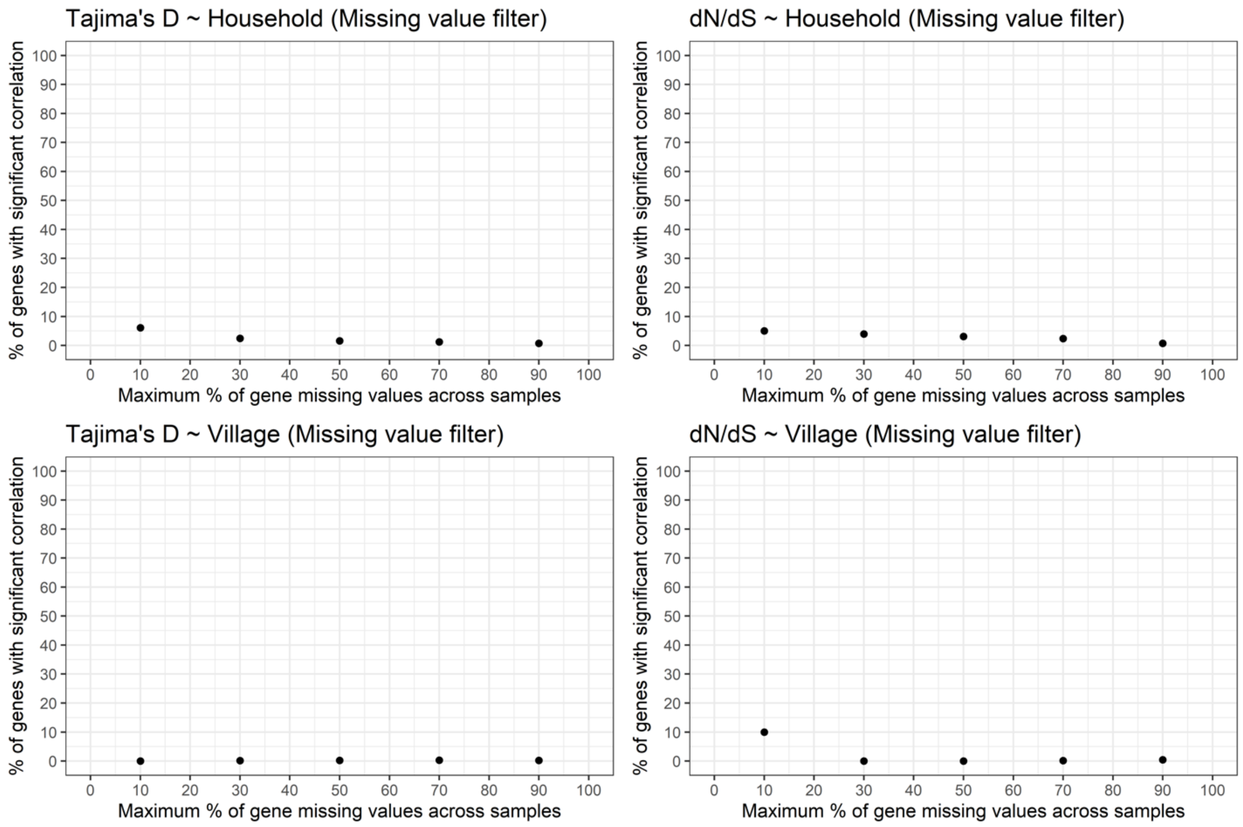


**Figure S5. The lack of significant correlations between host factors and mobile gene evolution is robust to filters imposed on missing data.**

This figure illustrates the percentage of genes for which the correlations between population genetics metrics and host factors are significant depending on the missing value filter stringency. The missing value filter defines a maximum percentage of samples in which the gene is absent or sequenced with less than 10X coverage. Increasing this threshold should increase the number of tested genes. **Figure 3A** shows that, using a 30% missing value threshold, population genetics metrics do not significantly correlate with host factors, except for a minority of genes (<5% of tested genes). This figure shows that the non-significance of this correlation is robust to the missing value filters because no matter the threshold chosen, the proportion of genes for which the correlations is significant is still very small. Because *Tajima's D* captures information from both $\theta_\pi$ and $\theta_w$, their correlations with host factors were not included in this figure.

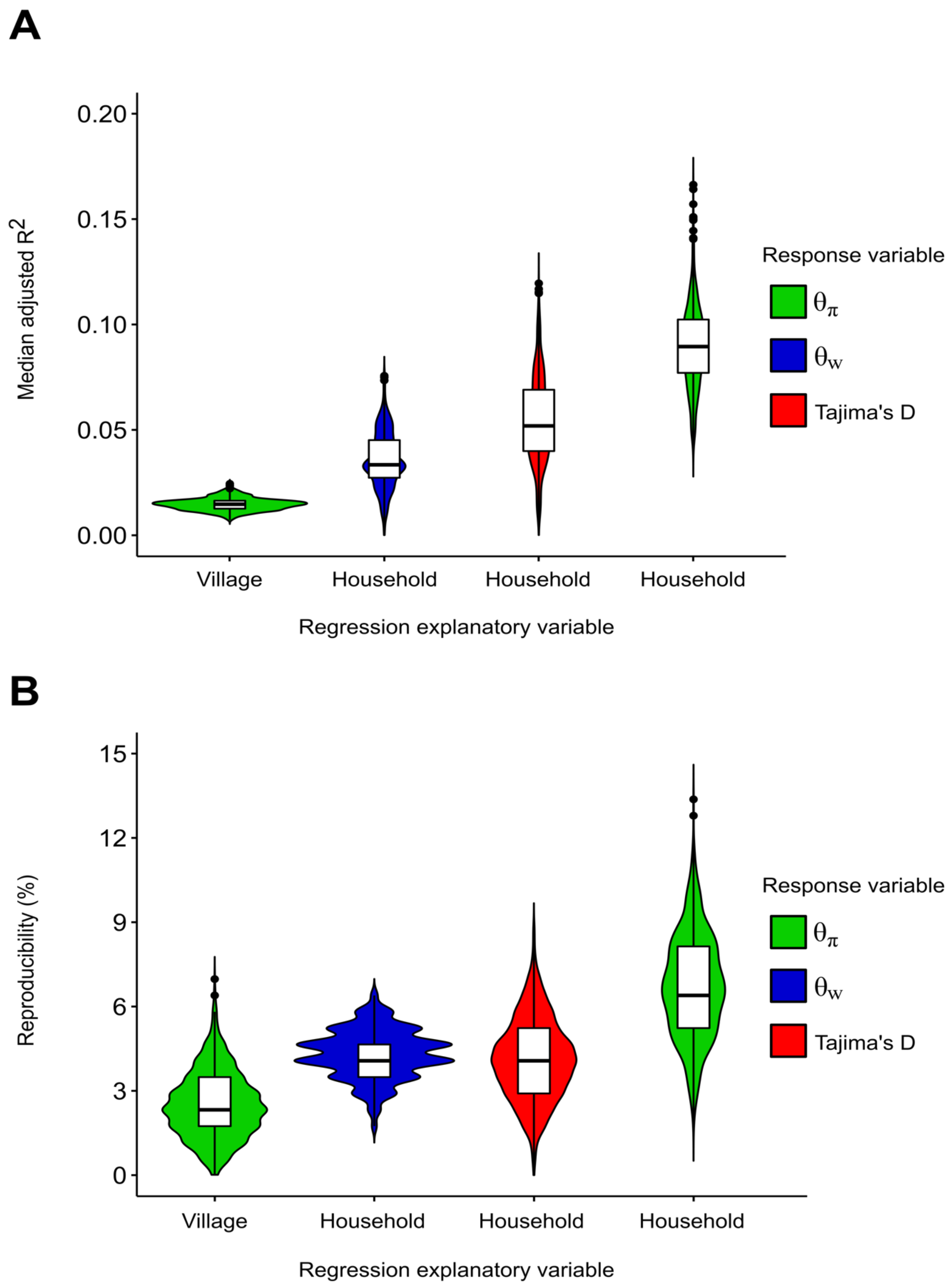


**Figure S6 Gene set size bias does not explain host attributes weak influence on mobile gene sequences evolution.** We hereby want to make sure that host attributes correlations with population genetics metrics were not weaker than COG family correlations only because their significance was assessed on a bigger set of objects (1333 genes vs 172 samples). We evaluated the significance of these correlations with the adjusted $R^2$ and the reproducibility, i.e. the proportion of regression objects (genes or samples) for which the regression is significant. Therefore, there could be a bias toward observing lower reproducibility in bigger set of objects. A) Median adjusted $R^2$ across subsamples. This figure shows the mean adjusted $R^2$ of top 4 regressions between population genetics metrics (color-coded) and host attributes across the 999 subsamples (n=172 genes). The downsampling do not increase the regression significance and rather supports the fact that it is weak because the median adjusted $R^2$ are low (<= 0.2). B) Reproducibility (%) across subsamples. Reproducibility represents the percentage of genes for which host attributes correlations are significant. The correlations and genes represented are the same than in **Figure S6A**. This figure shows that the reproducibility of these correlations, which ranges between 0 and 15%, is lower than the reproducibility of gene families correlations (**Figure 3B**), even after downsampling.

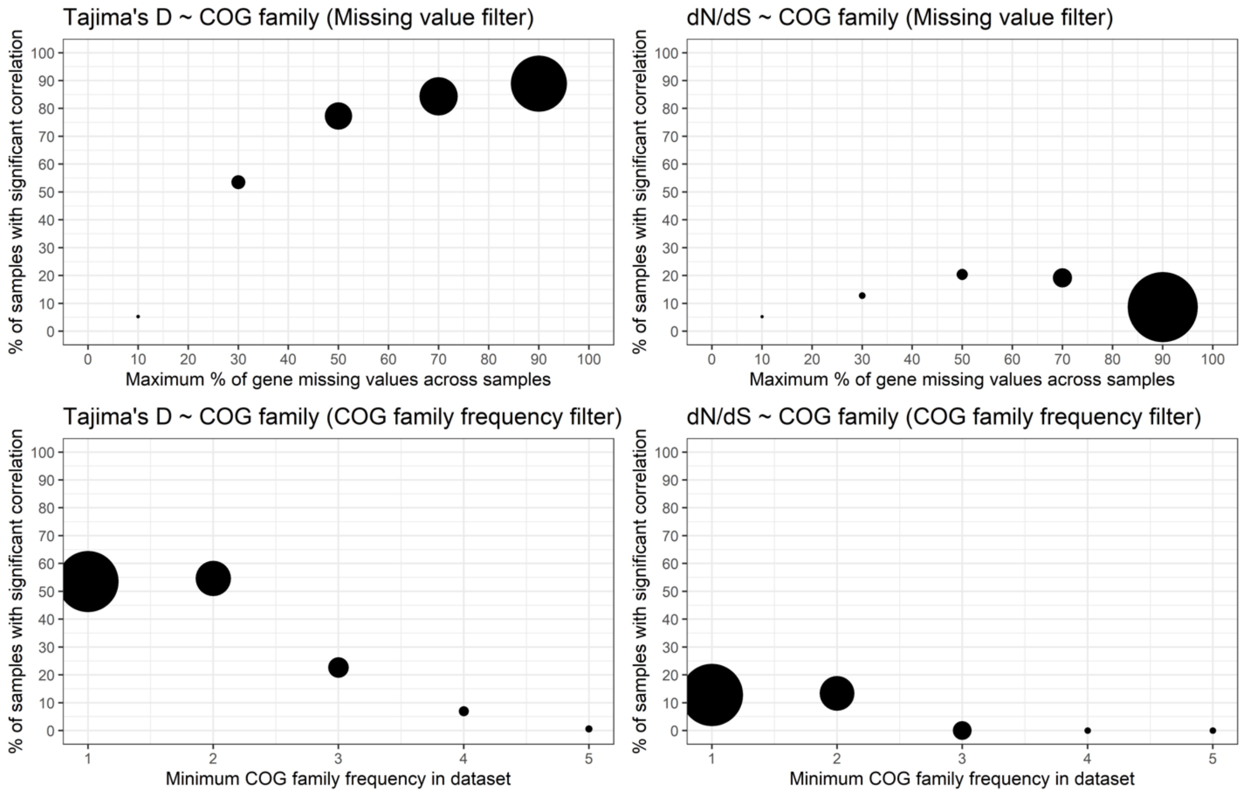


**Figure S7. COG family significant correlation with mobile genes evolution is not clearly generalizable to the whole dataset**

This figure illustrates the percentage of genes for which the correlations between population genetics metrics and COG family are significant depending on the missing value filter ($1^{st}$ row of the figure) and a filter on the minimum number of genes representing the COG family in the dataset ($2^{nd}$ row of the figure). The size of the points represents the relative sample size used for the correlations (normalized number of genes that passed the filters). The missing value filter defines a maximum percentage of samples in which the gene is absent or sequenced with less than 10X coverage. Thus, increasing this threshold makes this filter decrease in stringency and thus increase sample size. As for the filter on the minimum size of a COG family within the dataset, it defines a minimum threshold such that increasing it would increase stringency and thus decrease sample size. These filters have been chosen respectively to handle missing values caused by gene absence across sample (**Figure S12**) or gene with low coverage in gut metagenomes and to avoid the random effects of small sample size for COG families that are underrepresented in the dataset. Changing these thresholds should increase or decrease the number of tested genes. **Figure 3B** shows that, using a 30% missing value threshold and COG families with at least 2 genes in the dataset, population genetics metrics significantly correlate with COG family in the small subset of 512 tested genes. This figure shows that increasing the missing value filter stringency increases the strength of these correlations while the opposite is observed for the COG family minimum size filter due to reduced sample size. Because there seems to be a trade-off between the stringency of these filters and the amount of data used for the correlations, the strength of these correlations can fluctuate depending the quantity of tested genes. Therefore, we conclude that is not clear to which extent selective pressures on gene families are significant across the whole dataset (maximum sample size possible).

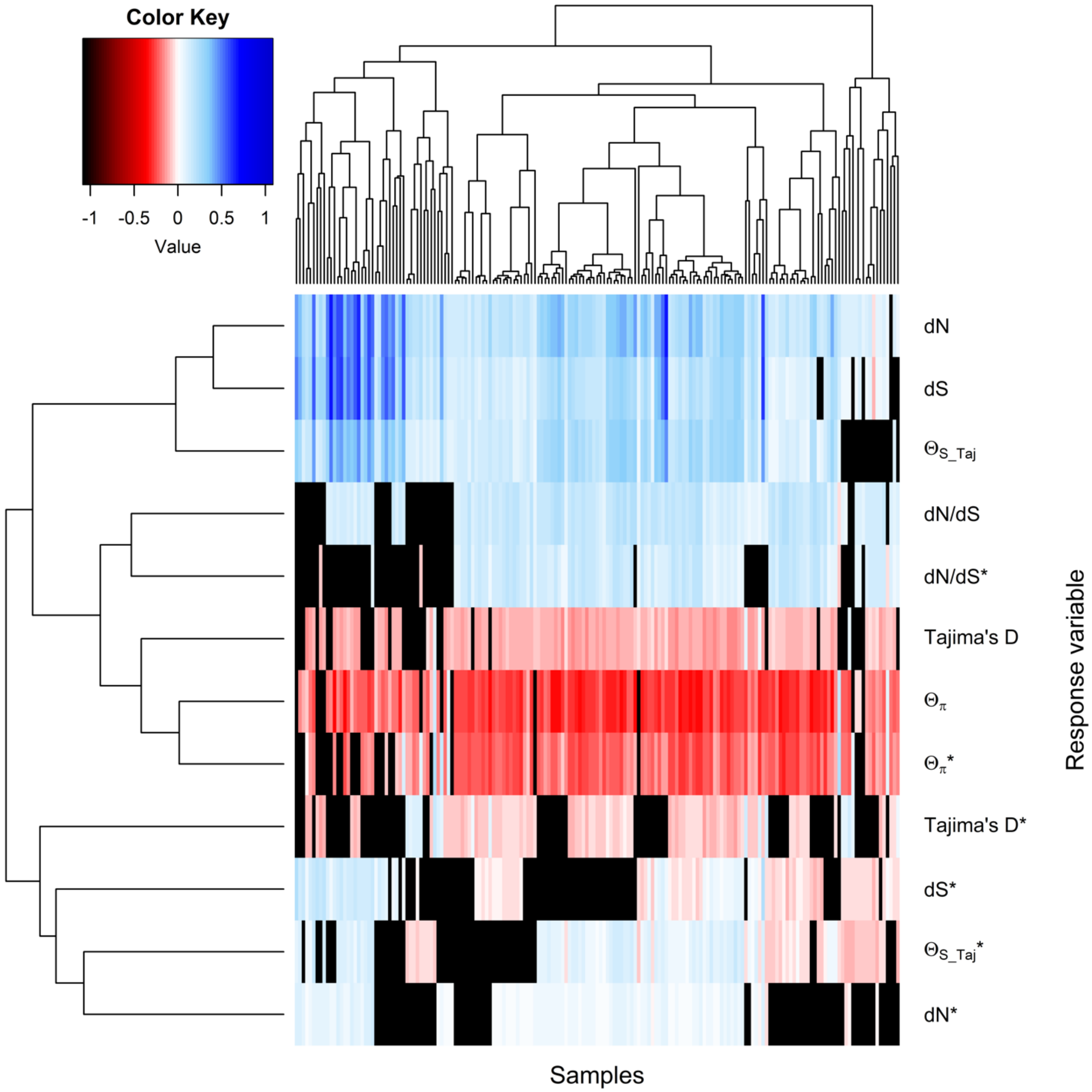


**Figure S8. Complete heatmap of gene mobility regression coefficients**

As mentioned in the result section 3, gene mobility has a significant impact on nucleotide diversity. We did two regression models for each response variable Y tested ($\theta_\pi$, $\theta_w$, *dN/dS* and *Tajima's D*): "Y ~ Gene Mobility" and "Y $*$ ~ Gene Mobility + Coverage + Gene length". The second type of model allows us to see the correlation between gene mobility and Y by controlling for coverage and gene length and thus sequencing artefacts like sequencing errors that increase with coverage and gene length. The effects of gene mobility on nucleotide diversity and *Tajima's D* were the only one supported in simulations (see the results section 3). Controlling for sequencing artefacts does not change these trends. We also observed a significant positive correlation between *dN/dS* and gene mobility across 144 out of the 175 samples (Bonferroni-adjusted p-value < 0.05). Nevertheless, this correlation was not supported in simulations and the adjusted $R^2$ was low, i.e. <10% for all samples.

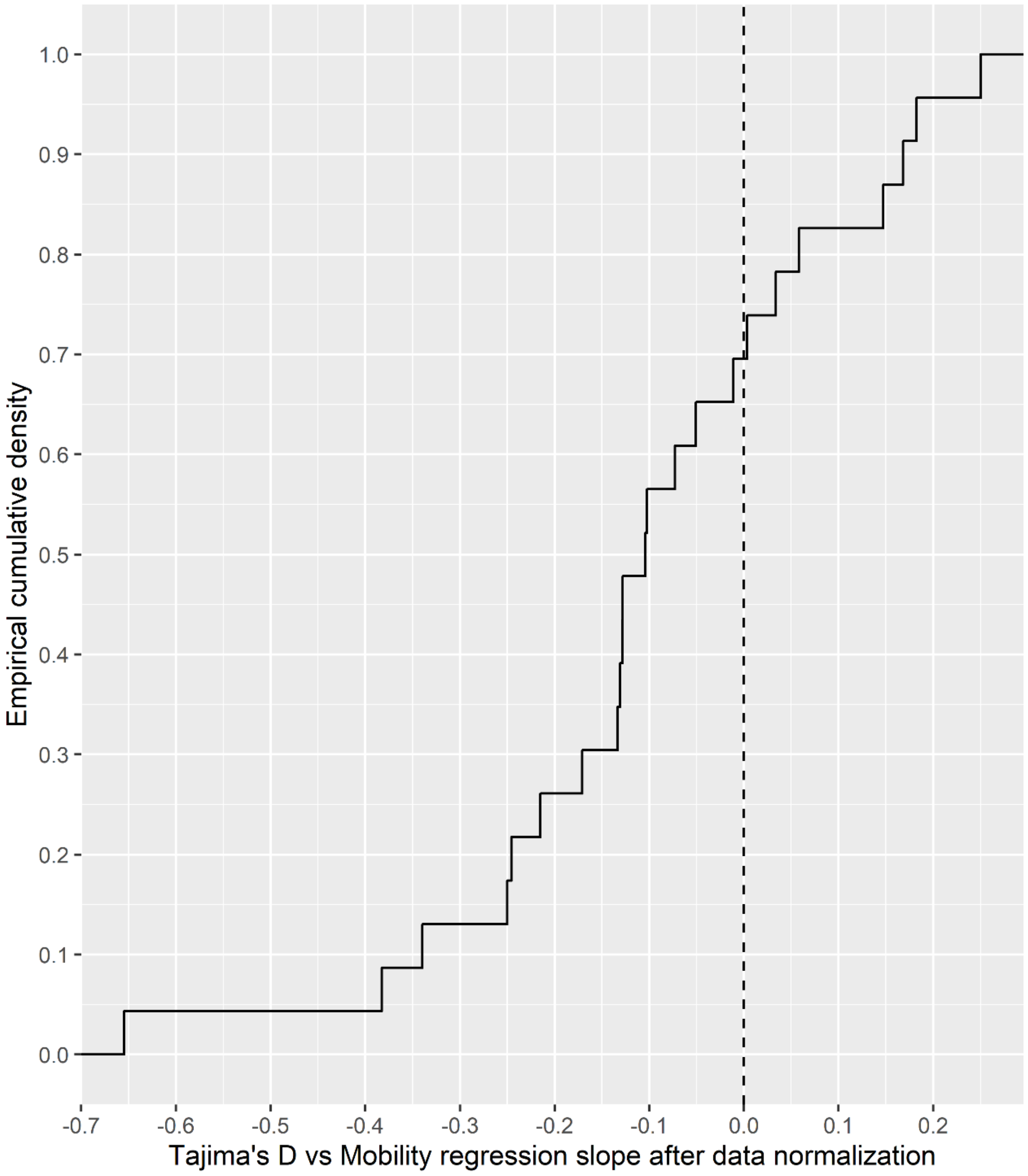


**Figure S9. Cumulative density distribution of "Tajima′s D ~ Mobility" regression slope across COG categories**

Data were normalized to make sure that regression residuals normality condition was respected before building the linear mixed model (*Tajima's D* Box-Cox $\lambda = 2$; Gene mobility Box-Cox $\lambda = -0.004658$). 3608 unique mobile genes data across the 175 samples were selected for this analysis. These mobile genes needed to pass the filters for variant analysis, i.e. mean site depth was >= 10, and have available COG annotations in the Metadata (Brito *et al.*, 2016). COG categories, for which *Tajima's D* is positively correlated to mobility, which is a deviation from the general trend, represent 30% of the distribution. As mentioned in section 3, these genes are part of COG categories that are related to important beneficial functions in the gut microbiome, i.e. P (Inorganic ion transport and metabolism), I (Lipid transport and metabolism), V (Defense mechanisms), Q (Secondary metabolites biosynthesis, transport and catabolism), U(Intracellular trafficking, secretion, and vesicular transport), O(Posttranslational modification, protein turnover, chaperones) and G(Carbohydrates transport and metabolism).

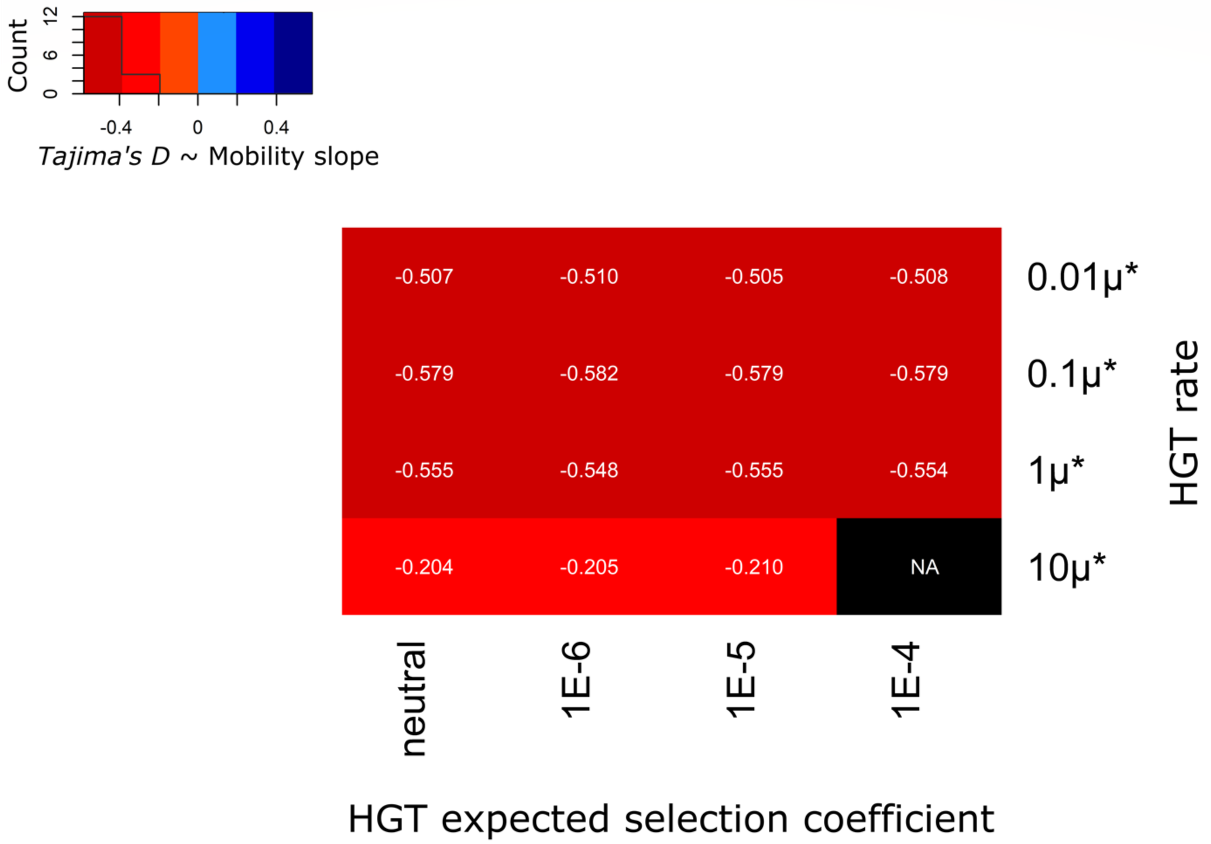


**Figure S10. Mobile genes nucleotide diversity is more influenced by HGT rate than HGT selection coefficient in simulations.** This figure shows the heatmap of "$Tajima's\ D \sim$ Gene Mobility" average slope across simulations. Regression p-values were obtained through a t-test. Data standardization was performed before each regression to respect t-test assumption of normality. The slope is indicated in white in the heatmap cells and its sign is represented by the color, i.e. red represents a negative slope, blue represents a positive slope and black represent non-significant results (NA). Across simulations, the slope of the regression varies more because of HGT rate changes than HGT fitness effect. This was confirmed by a K-S test (p-value < 0.05). The test was realized with 160 simulations data and the average slopes were calculated over the 10 replicates of each of the 16 simulation sets (Only the significant slopes were considered). Simulation sets differ by HGT rate or HGT fitness effect: HGT rate $\in$ $[0.01\mu^*, 0.1\mu^*, 1\mu^*, 10\mu^*]$ and HGT $\lambda \in$ $[\text{neutral}, 10^4, 10^5, 10^6]$, where $\lambda$ is the rate parameter of the exponential distribution of HGT selection coefficient and $\mu^*$ is the mutation rate (Methods). The expected selection coefficient corresponds to $\frac{1}{\lambda}$. We increased Prokaryotic mutation rate $\mu^*$ up to the order of $10^{-7}$ mutations per site per generation for practical reason and made sure it does not change genome size equilibrium (Methods and **Figures S12**). Each simulation included 5000 cells, 10 species, 500 genes per cells at equilibrium and start, as well as a simulation time of $10^5$ generations.

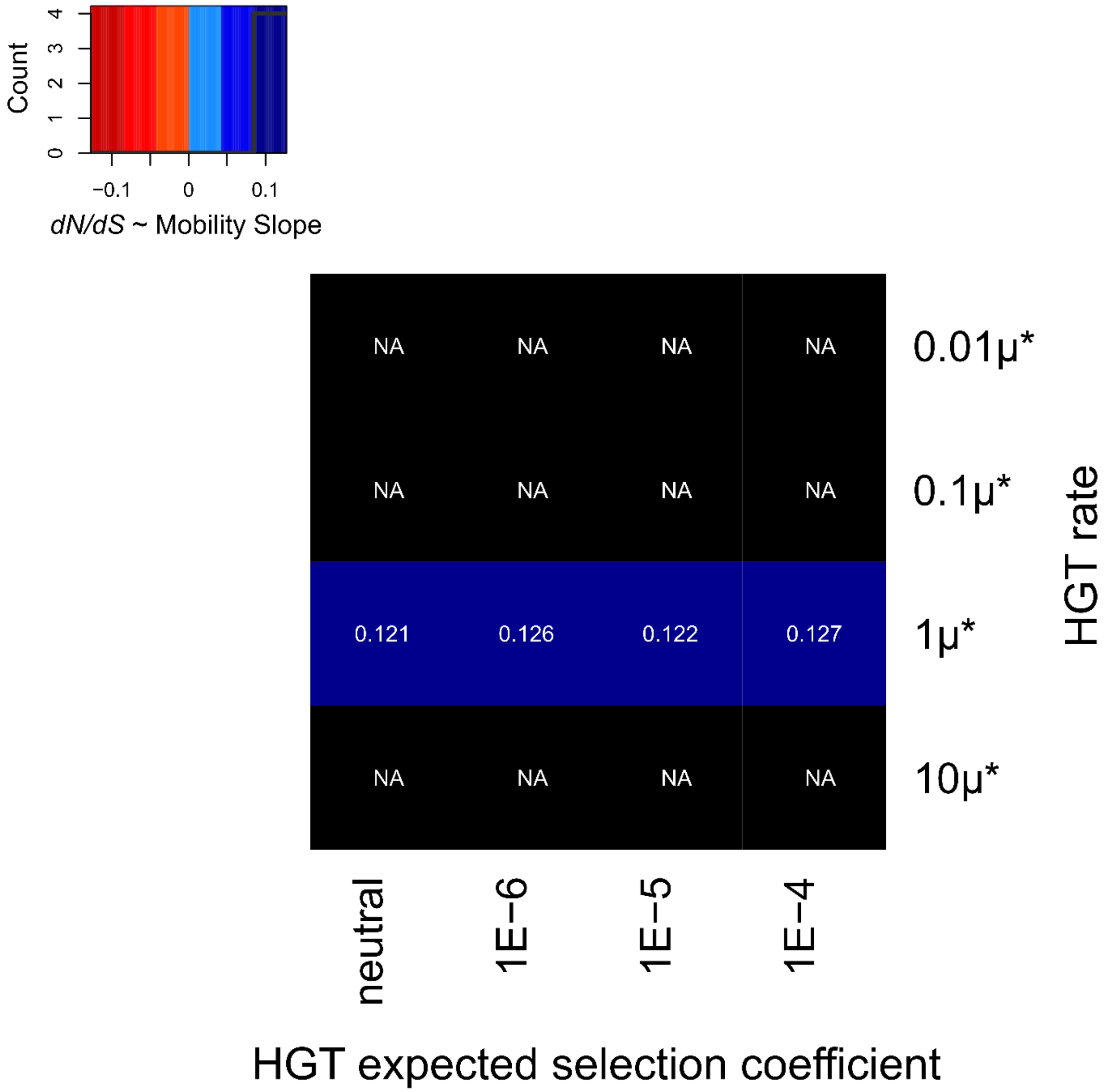


**Figure S11. *dN/dS* correlates weakly with gene mobility in simulations.**

Heatmap of "dN/dS ~ Gene Mobility" average slope across simulations Regression p-values were obtained through a t-test. Data standardization was performed before each regression to respect t-test assumption of normality. The slope is indicated in white in the heatmap cells and its sign is represented by the color, i.e. red represents a negative slope, blue represents a positive slope and black represent non-significant results (NA). In the simulation for which the correlation is significant, the results show that *dN/dS* tends to be positively correlated with Gene mobility. However, the correlation is not reproducible across most of the simulations and the average adjusted $R^2$ was 0.01, which is very weak. The simulations included 5000 cells, 10 species, 500 genes per cells at equilibrium and simulation time of $10^5$ generations.

**Figures S12. Genome size equilibrium across simulations**

**(See Figure_S12_Genome_Size_Equilibrium_All_Simulations.pdf)**

This set of figures represents the time series of genome size during the simulations mentioned in this manuscript: a) HGT rate = 0.01μ and HGT is neutral, b) HGT rate = 0.01μ and λ = 1E6, c) HGT rate = 0.01μ and λ = 1E5, d) HGT rate = 0.01μ and λ = 1E4, e) HGT rate = 0.1μ and HGT is neutral, f) HGT rate = 0.1μ and λ = 1E6, g) HGT rate = 0.1μ and λ = 1E5, h) HGT rate = 0.1μ and λ = 1E4, i) HGT rate = 1μ and HGT is neutral, j) HGT rate = 1μ and λ = 1E6, k) HGT rate = 1μ and λ = 1E5, l) HGT rate = 0.01μ and λ = 1E4, m) HGT rate = 10μ and HGT is neutral, n) HGT rate = 10μ and λ = 1E6, o) HGT rate = 10μ and λ = 1E5 and p) HGT rate = 10μ and λ = 1E4, where the expected selection coefficient of HGT (*s*) = $\frac{1}{\lambda}$. It is important to show that these time series are on dynamic equilibrium, i.e. genome size fluctuates around a certain value, because it supports the fact that our results are not dependent on the initial conditions of the simulations.

**Tables S1 Comparison of full and nested linear mixed models**

| Model | Description | $R^2$ | Log-likelihood | LRT p-value (Nested model vs M1 model) |
|---|---|---|---|---|
| M1 | Coverage vs Mobility + all random factors, i.e. Sample and COG category | 0.35 | -317537 | - |
| M2 | Coverage vs Mobility + COG category (without Sample) | 0.12 | -355586 | $< 2.2x10^{-16}$ |
| M3 | Coverage vs Mobility + Sample (without COG category) | 0.31 | -321438 | $< 2.2x10^{-16}$ |

**Table S1A Regression strength of the linear mixed model "Coverage ~ Mobility + Sample + COG category" and nested models LRT.** Because the likelihood ratio test (LRT) p-value is highly significant for each of the nested models, all the random effects tested, i.e. sample and COG category, have a significant effect on the regression. The M3 model $R^2$ is much higher than the M2 model $R^2$, suggesting that the relationship between Coverage and Mobility varies more across samples than across COG categories.

| Model | Description | $R^2$ | Log-likelihood | LRT p-value (Nested model vs M1 model) |
|---|---|---|---|---|
| M1 | *Tajima's D* vs Mobility + all random factors, i.e. Sample and COG category | 0.07 | -454557 | - |
| M2 | *Tajima's D* vs Mobility + COG category (without Sample) | 0.04 | -457645 | $< 2.2x10^{-16}$ |
| M3 | *Tajima's D* vs Mobility + Sample (without COG category) | 0.033 | -458084 | $< 2.2x10^{-16}$ |

**Table S1B Regression strength of the linear mixed model "Tajima's D ~ Mobility + Sample + COG category" and nested models LRT.** Because the likelihood ratio test (LRT) p-value is highly significant for each of the nested models, all the random effects tested, i.e. sample and COG category, have a significant effect on the regression. M2 model $R^2$ is much higher than M3 model $R^2$, suggesting that the relationship between *Tajima's D* and Mobility varies more across COG categories than across samples. The $R^2$ of this linear mixed model, i.e. "$Tajima's\ \mathrm{D} \sim \mathrm{Mobility} + \mathrm{Sample} + \mathrm{COG\ category}$", is much lower than the $R^2$ of the mixed model "$\mathrm{Coverage} \sim \mathrm{Mobility} + \mathrm{Sample} + \mathrm{COG\ category}$" (**Table S1A**), because mobility correlates negatively with *Tajima's D* for some COG categories and positively for others (**Figure 5A**) so that the general trend is a weak, but significant negative correlation, contrarily to Coverage which always correlates positively with mobility (**Figure 1**).

| Model | Description | $R^2$ | Log-likelihood | LRT p-value (Nested model vs M1 model) |
|---|---|---|---|---|
| M1 | FPKM vs Mobility + all random factors, i.e. Sample and COG category | 0.321 | -3346029 | - |
| M2 | FPKM vs Mobility + COG category (without Sample) | 0.205 | -3456259 | $< 2.2 \times 10^{-16}$ |
| M3 | FPKM vs Mobility + Sample (without COG category) | 0.284 | -3362567 | $< 2.2 \times 10^{-16}$ |

**Table S1C Regression strength of the linear mixed model "$\mathbf{FPKM \sim Mobility + Sample + COG\ category}$" and nested models LRT.** FPKM represents the number of fragments that mapped to a gene sequence normalized by the length of the gene in kilobase and the number of reads produced from the sample in million of mapped reads. It is a common metric of gene relative abundance and it is positively correlated with mobility (p-value = 9.17E-07 < 0.05). Here we show that all the random effects tested, i.e. sample and COG category, have a significant effect on the regression because the likelihood ratio test (LRT) p-value is highly significant for each of the nested models. The M3 model $R^2$ is much higher than the M2 model $R^2$, suggesting that the relationship between FPKM and Mobility varies more across samples than across COG categories, as it is the case for Coverage.

**Tables S2 Metadata about mobile genes for population genetics metrics significantly correlate with host household or village.** These tables contain the information about A) genes for which *dN/dS* significantly correlates with host household, B) genes for which *Tajima's D* significantly correlates with host household and C) genes for which $\theta_\pi$ significantly correlates with host Village. In table S1C, COG categories that are related to the subset of functions that Brito et al. (2016) identified as being particularly abundant or prevalent in certain Fiji villages are highlighted in yellow. The tables were extracted from publicly available metadata from Fiji Community Microbiome Project (http://fijicomp.bme.cornell.edu//data.html):

| Simulation (n=10 replicates) | Average slope (standard deviation σ) | Average adjusted $R^2$ (standard deviation σ) | Average p-value (with Bonferroni correction) |
|---|---|---|---|
| HGT rate = 0.1µ; HGT λ = 1E4 | 0.958 (σ = 0.002) | 0.917 (σ = 0.004) | < $2.2 \times 10^{-16}$ |
| HGT rate = 0.1µ; neutral HGT | 0.958 (σ = 0.002) | 0.917 (σ = 0.004) | < $2.2 \times 10^{-16}$ |
| HGT rate = 1µ; HGT λ = 1E4 | 0.855 (σ = 0.006) | 0.731 (σ = 0.011) | < $2.2 \times 10^{-16}$ |
| HGT rate = 1µ; neutral HGT | 0.860 (σ = 0.006) | 0.740 (σ = 0.010) | < $2.2 \times 10^{-16}$ |
| HGT rate = 10µ; HGT λ = 1E4 | 0.684 (σ = 0.238) | 0.467 (σ = 0.033) | < $2.2 \times 10^{-16}$ |
| HGT rate = 10µ; neutral HGT | 0.674 (σ = 0.251) | 0.454 (σ = 0.034) | < $2.2 \times 10^{-16}$ |

**Table S3 Simulations support the positive correlation between gene census population size and Mobility.** Gene census population size was estimated with the number of copies of the gene. Each simulation included 5000 cells, 10 species, 500 genes per cells at equilibrium and a simulation time of $10^5$ generations. No matter if HGT was neutral or adaptive, this positive correlation was strongly supported (average adjusted $R^2$ over the 60 simulation replicates = 0.705; standard deviation = 0.190; p-value < $2.2 \times 10^{-16}$ for all replicates).
