## Supplementary material for "Pangenome sequence evolution within human gut microbiomes is explained by gene-specific rather than host-specific selective pressures": Figure S12

**a**

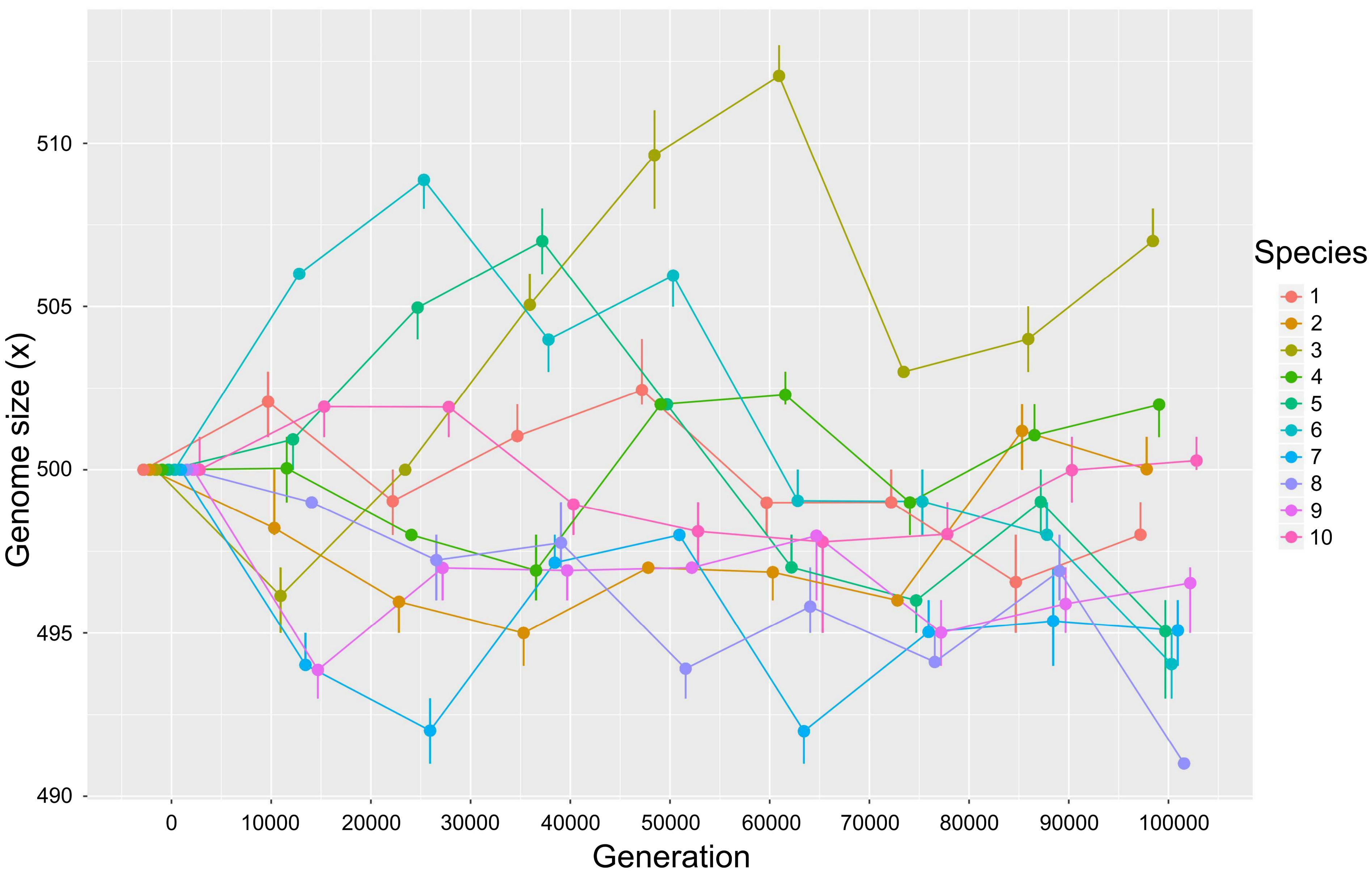
Genome size (x)
510
505
500
495
490
0
10000
20000
30000
40000
50000
60000
70000
80000
90000
100000
Generation
Species
1
2
3
4
5
6
7
8
9
10

**b**

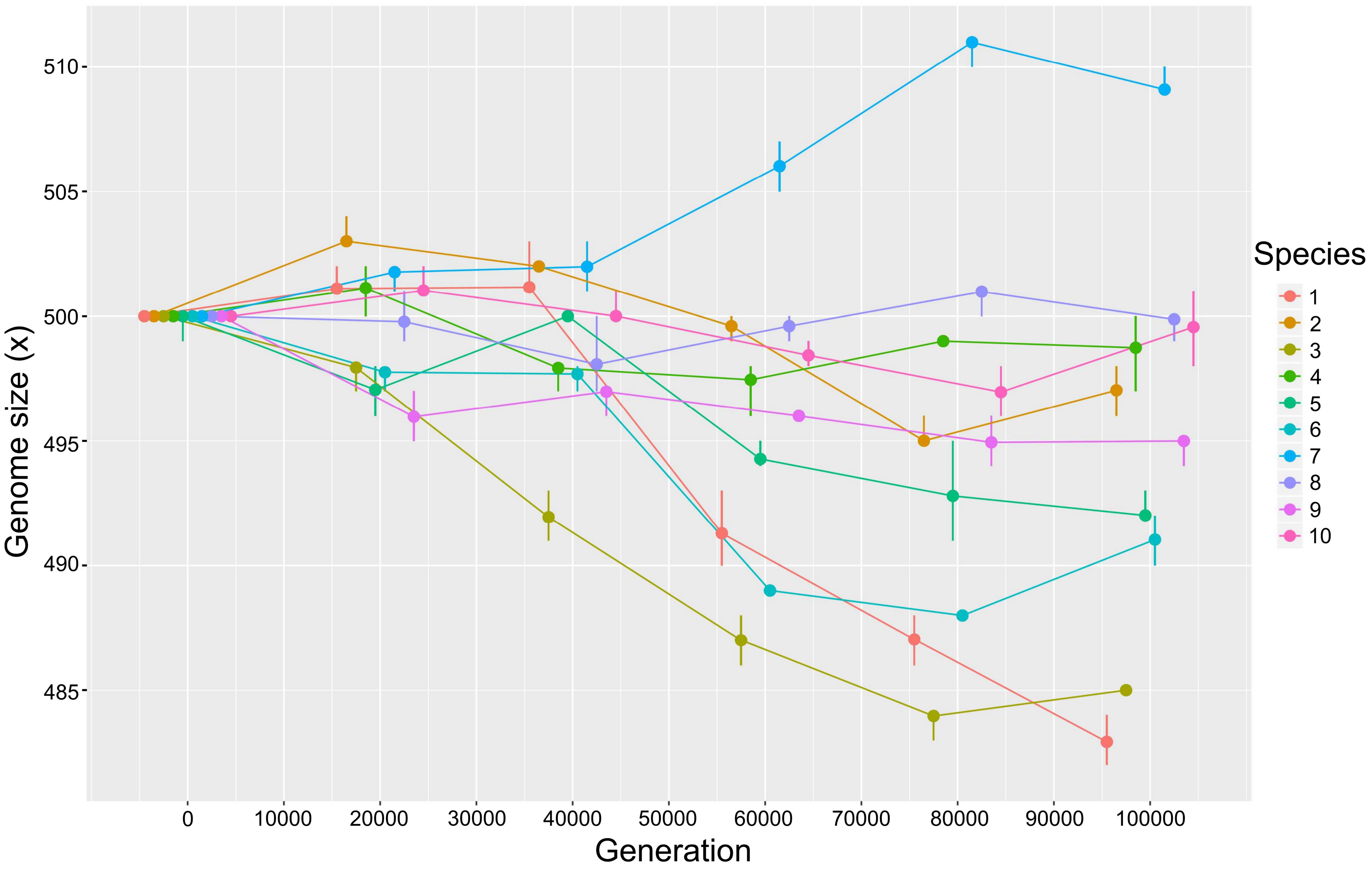

Genome size (x)
510
505
500
495
490
485
0
10000
20000
30000
40000
50000
60000
70000
80000
90000
100000
Generation
Species
1
2
3
4
5
6
7
8
9
10

**c**

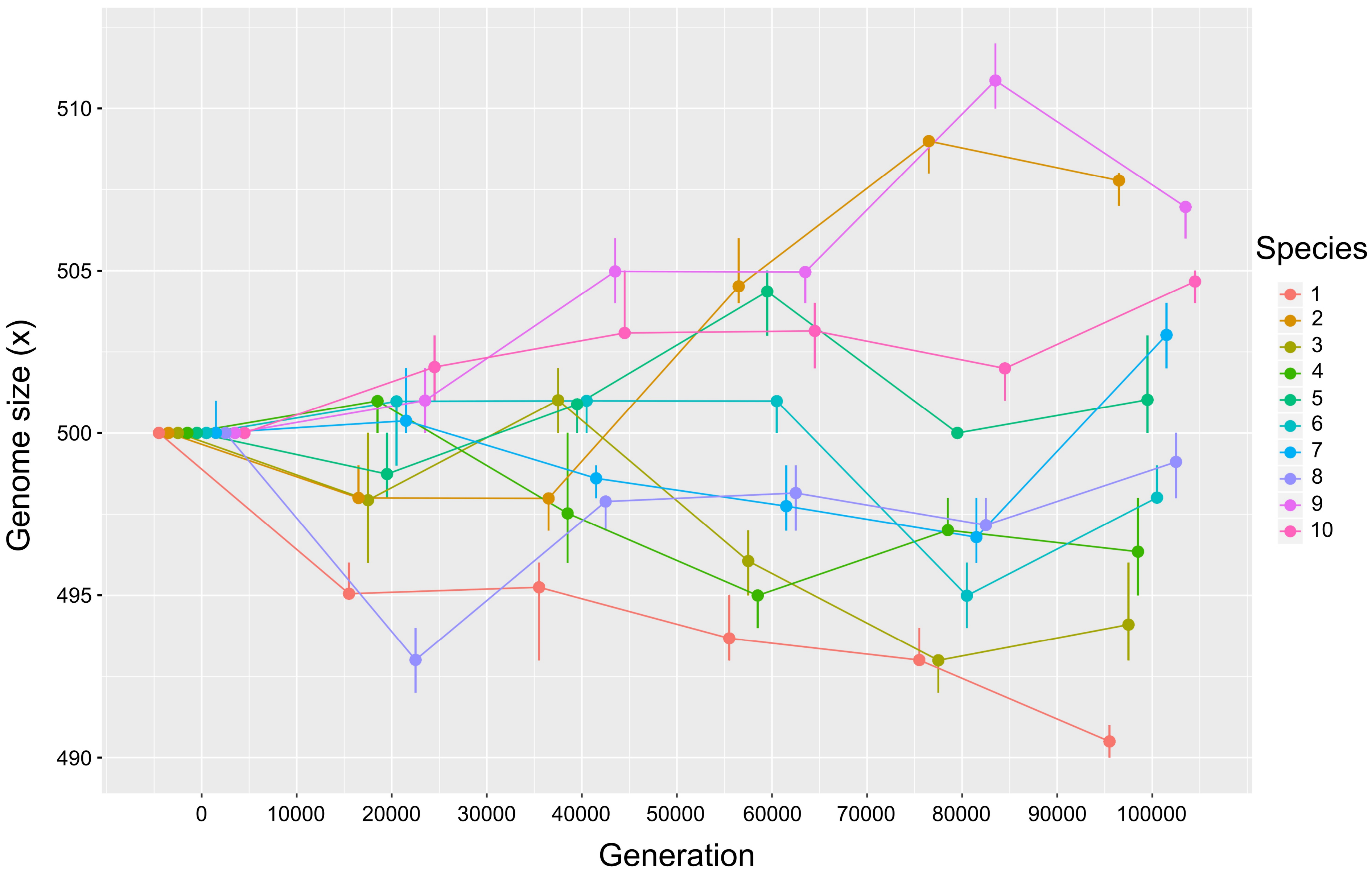

Genome size (x)
510
505
500
495
490
0
10000
20000
30000
40000
50000
60000
70000
80000
90000
100000
Generation
Species
1
2
3
4
5
6
7
8
9
10

d

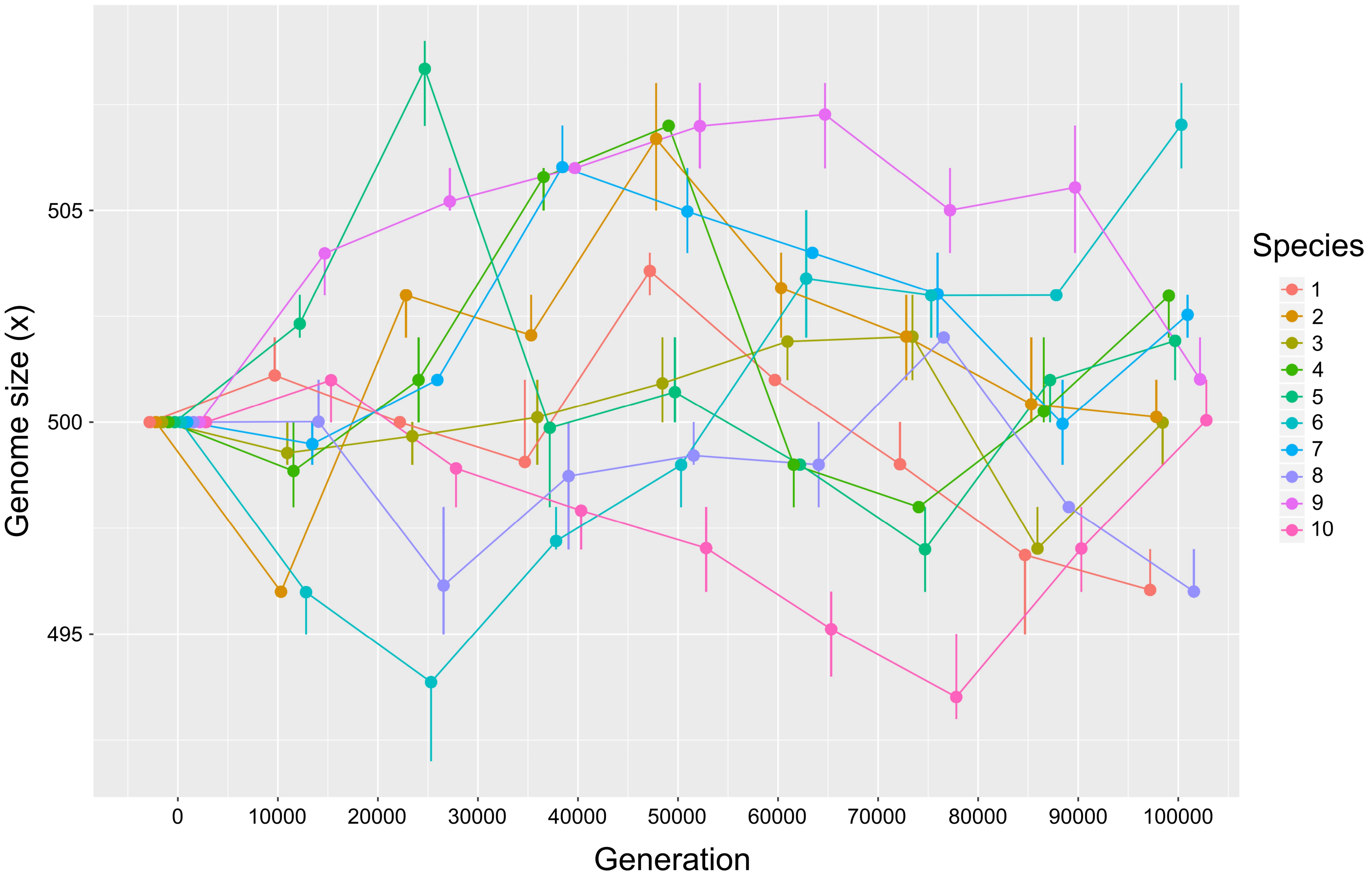
Genome size (x)
505
500
495
0
10000
20000
30000
40000
50000
60000
70000
80000
90000
100000
Generation
Species
1
2
3
4
5
6
7
8
9
10

**e**

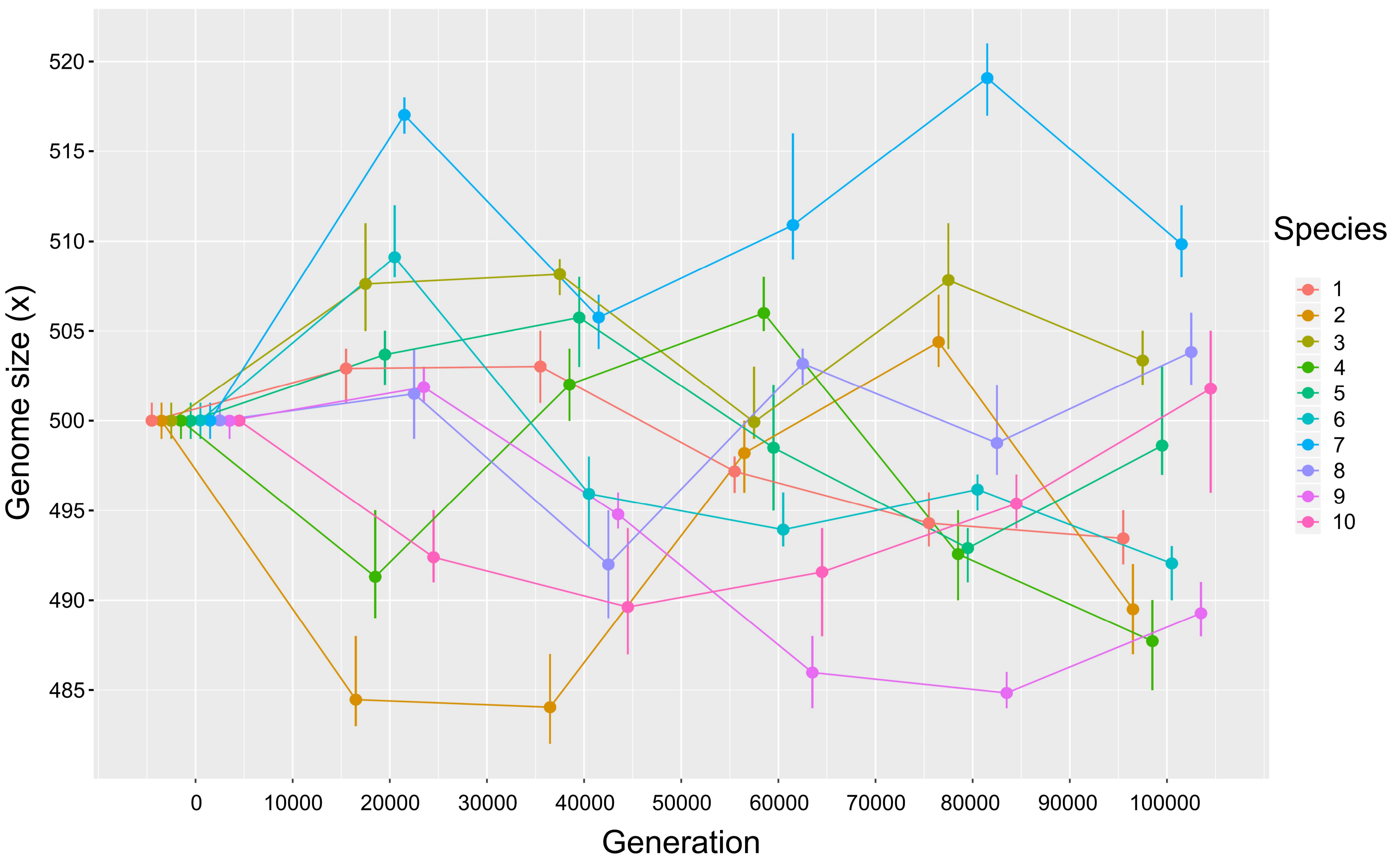

Genome size (x)
520
515
510
505
500
495
490
485
Generation
0
10000
20000
30000
40000
50000
60000
70000
80000
90000
100000
Species
1
2
3
4
5
6
7
8
9
10

f

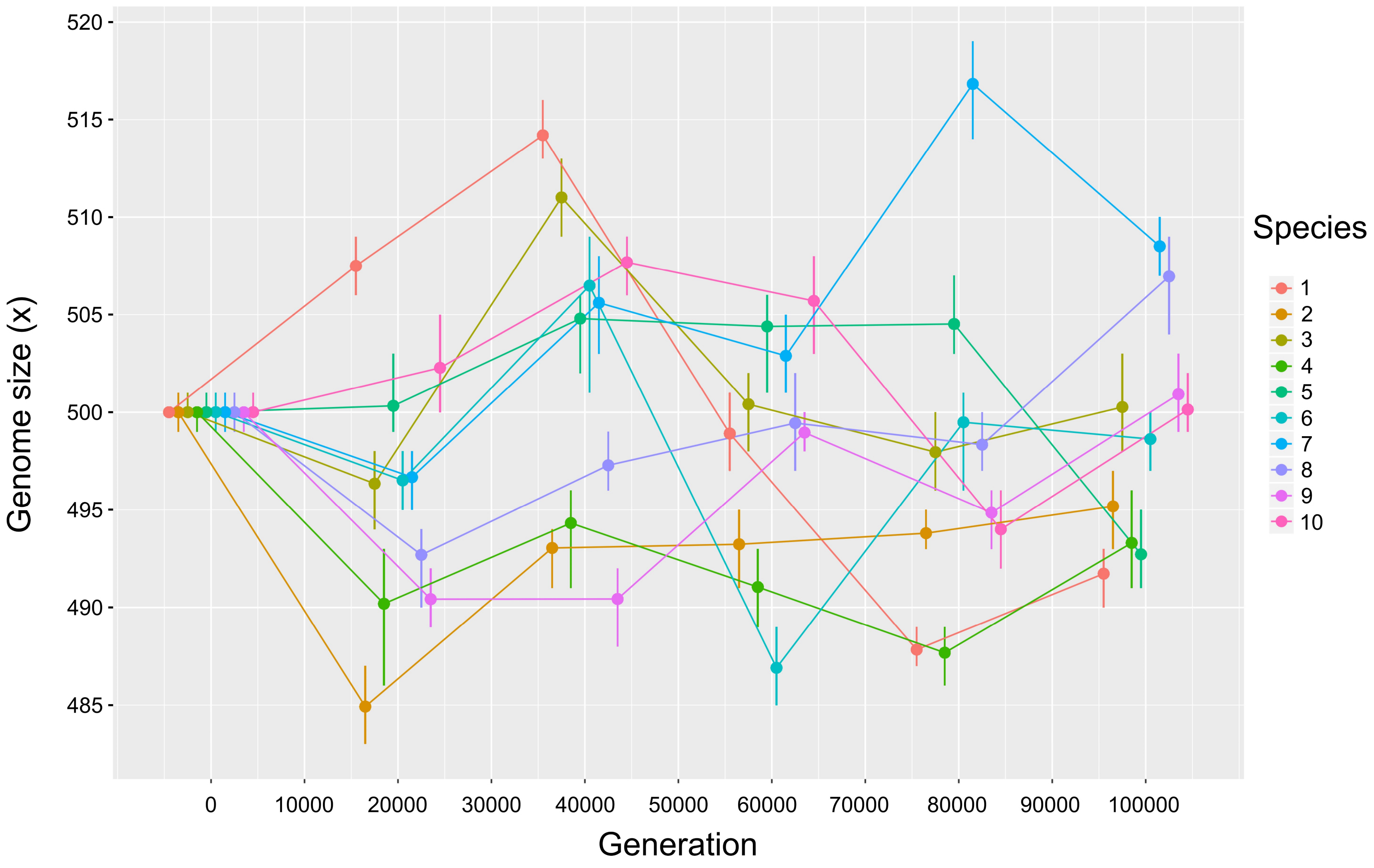
Genome size (x)
520
515
510
505
500
495
490
485
0
10000
20000
30000
40000
50000
60000
70000
80000
90000
100000
Generation
Species
1
2
3
4
5
6
7
8
9
10

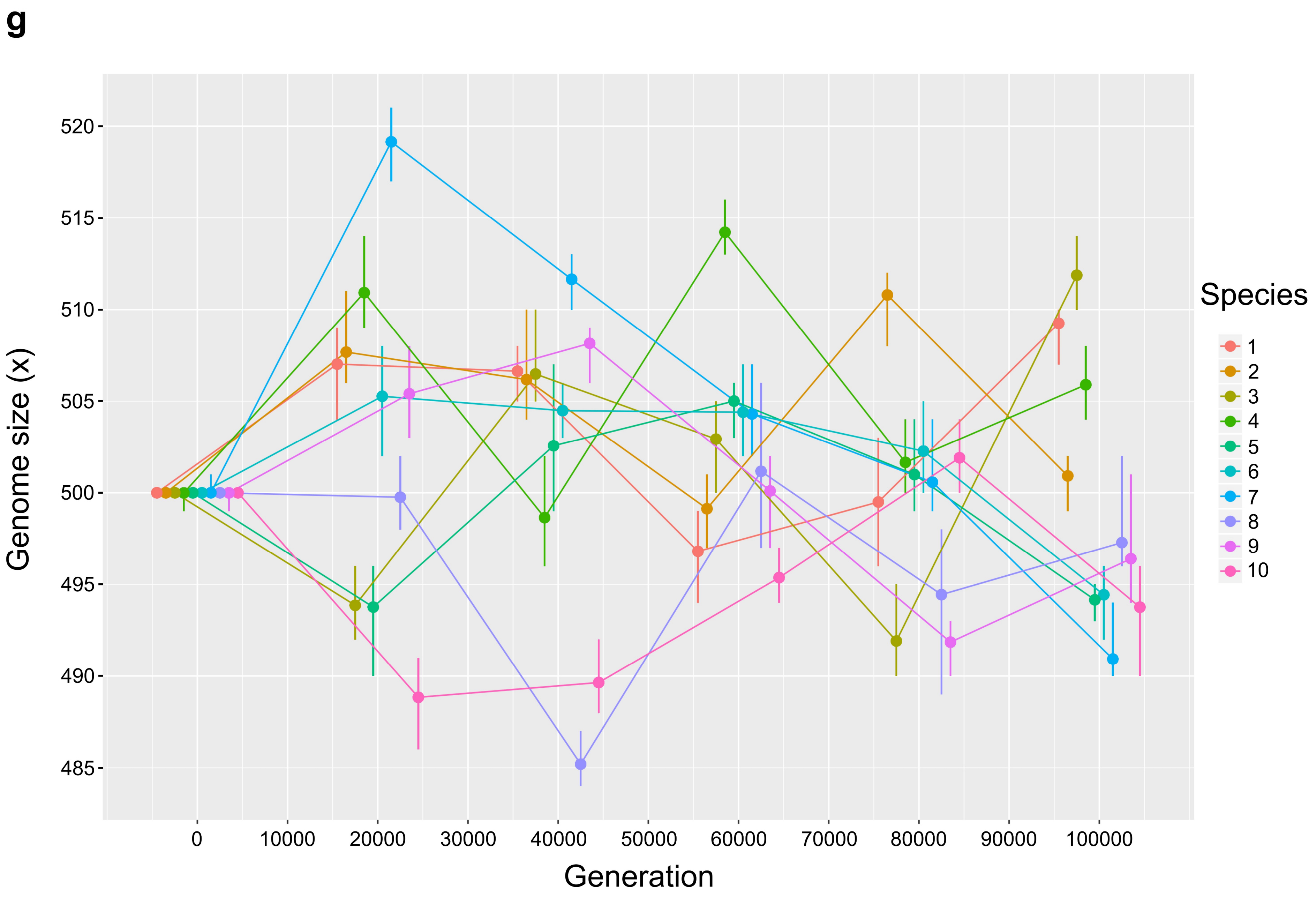
g
Genome size (x)
520
515
510
505
500
495
490
485
0
10000
20000
30000
40000
50000
60000
70000
80000
90000
100000
Generation
Species
1
2
3
4
5
6
7
8
9
10

**h**

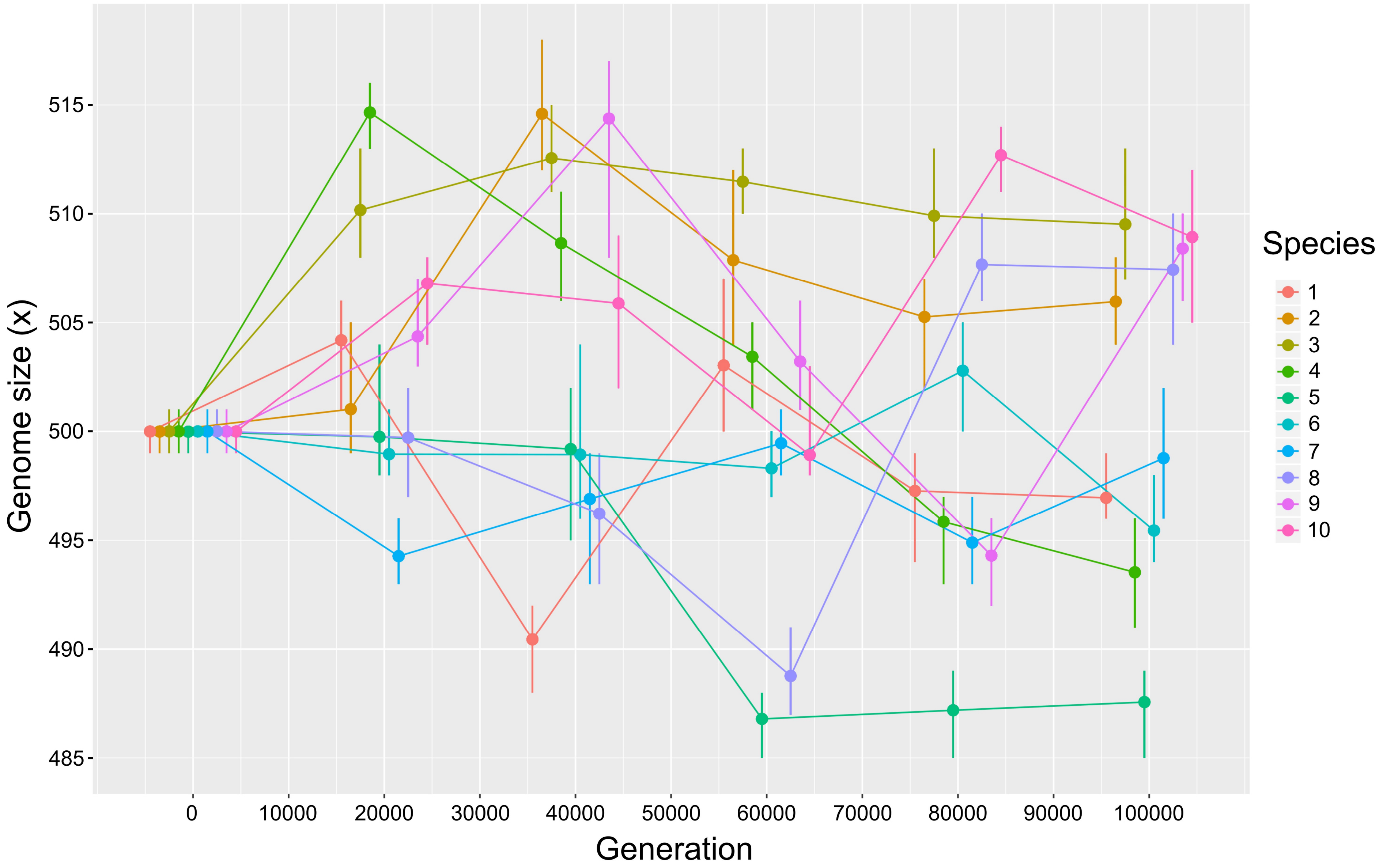

Genome size (x)
515
510
505
500
495
490
485
0
10000
20000
30000
40000
50000
60000
70000
80000
90000
100000
Generation
Species
1
2
3
4
5
6
7
8
9
10

**i**

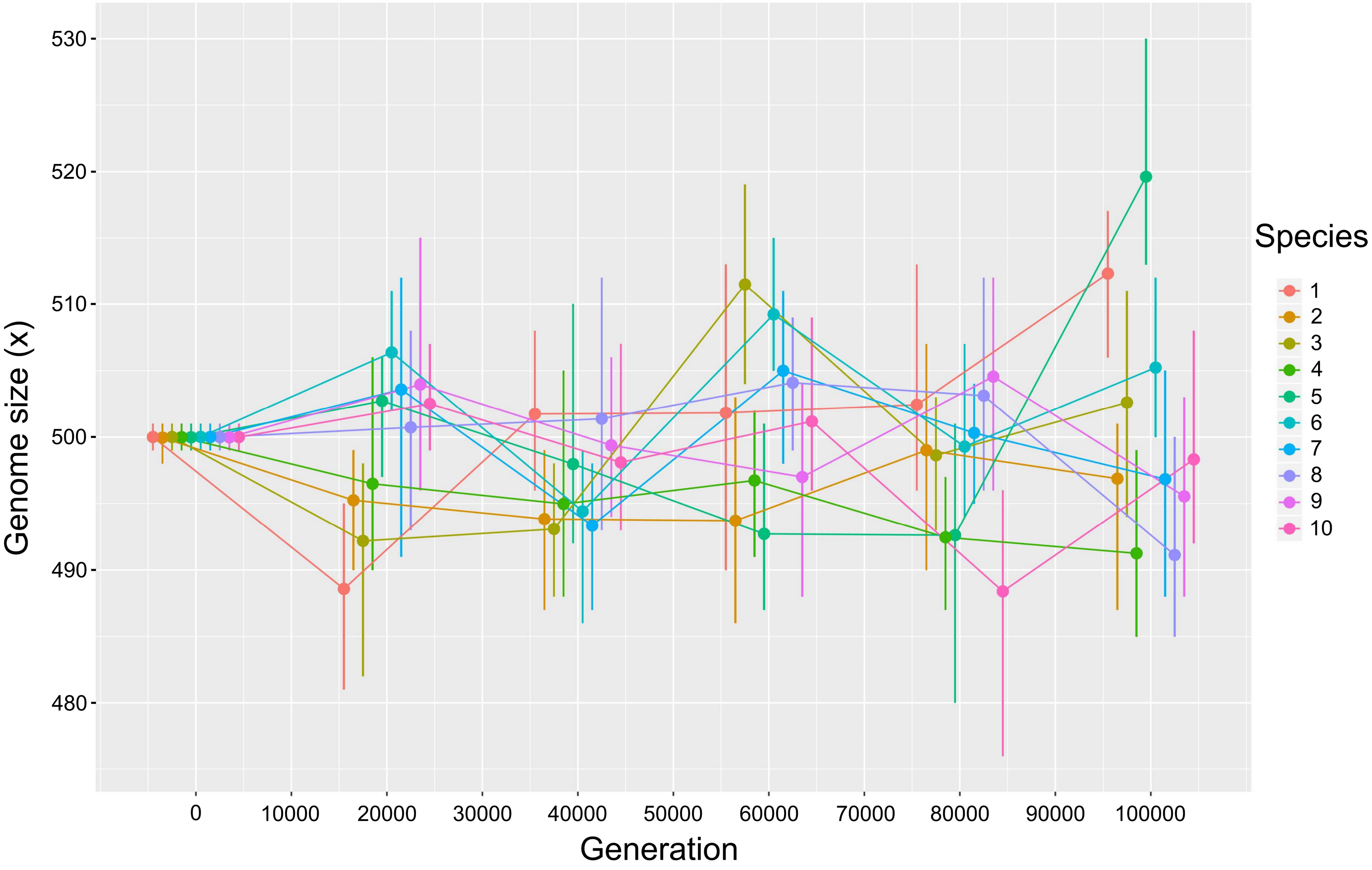

Genome size (x)
530
520
510
500
490
480
0
10000
20000
30000
40000
50000
60000
70000
80000
90000
100000
Generation
Species
1
2
3
4
5
6
7
8
9
10

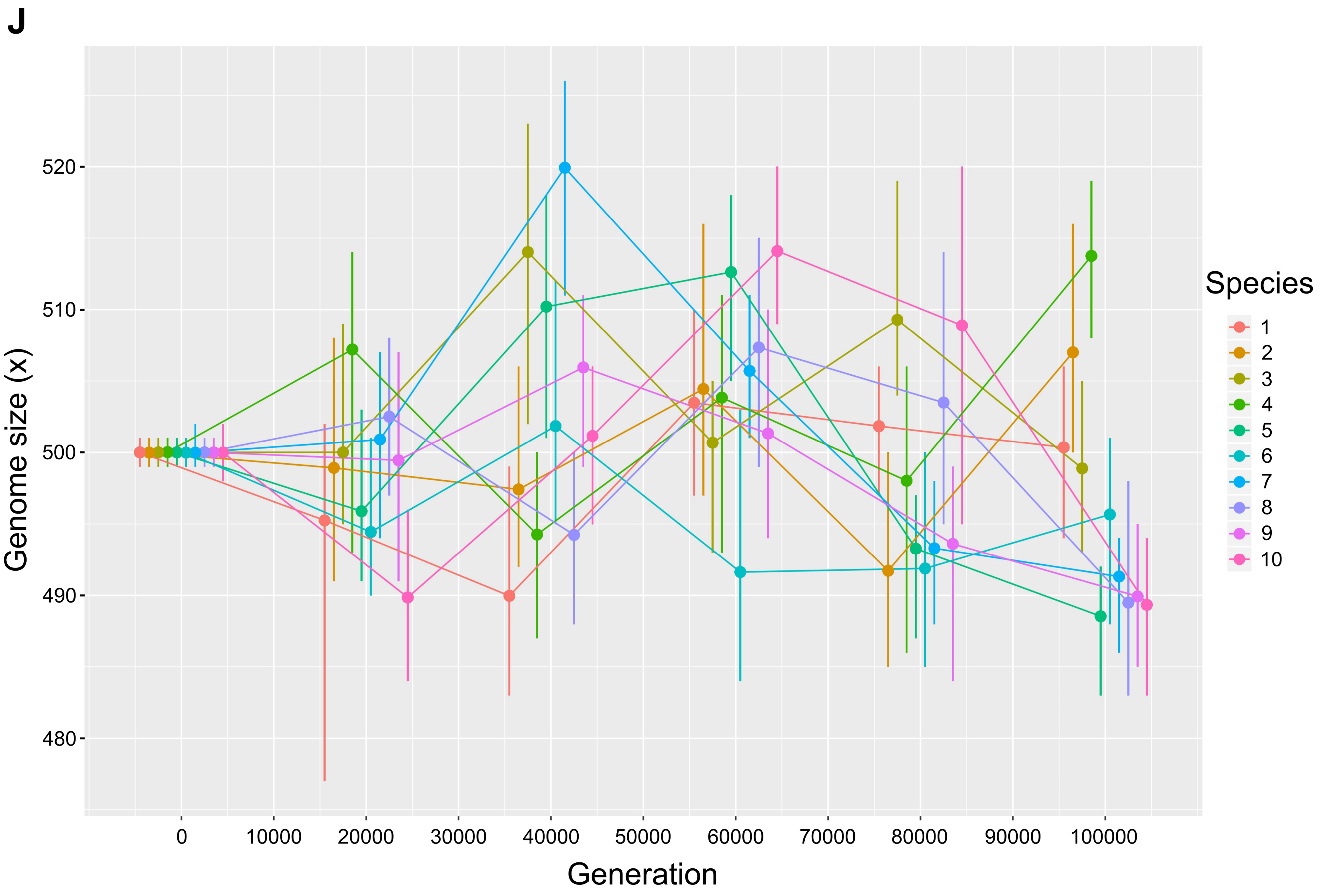

J
Genome size (x)
520
510
500
490
480
0
10000
20000
30000
40000
50000
60000
70000
80000
90000
100000
Generation
Species
1
2
3
4
5
6
7
8
9
10

**k**

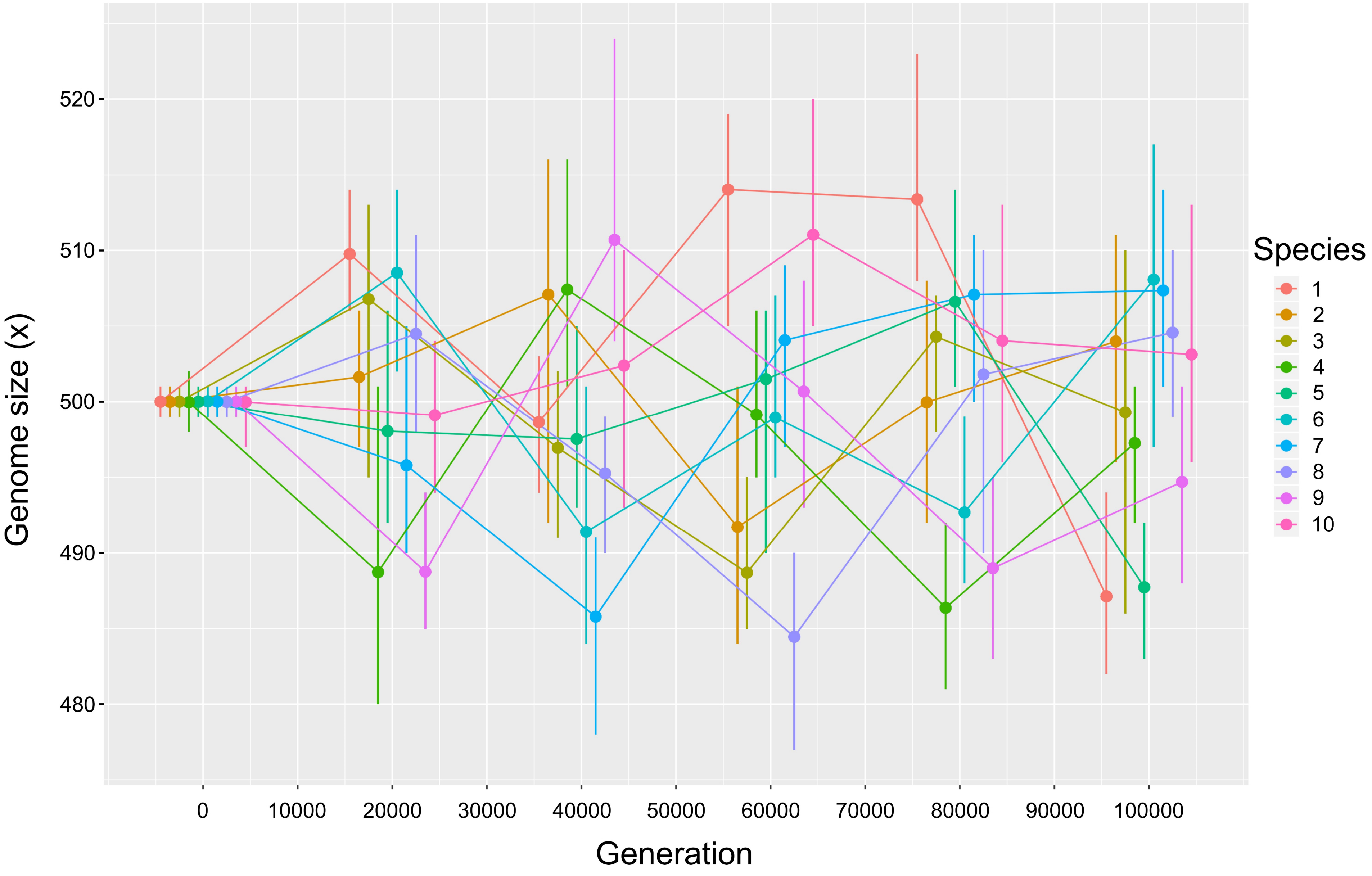
Genome size (x)
520
510
500
490
480
0
10000
20000
30000
40000
50000
60000
70000
80000
90000
100000
Generation
Species
1
2
3
4
5
6
7
8
9
10

I

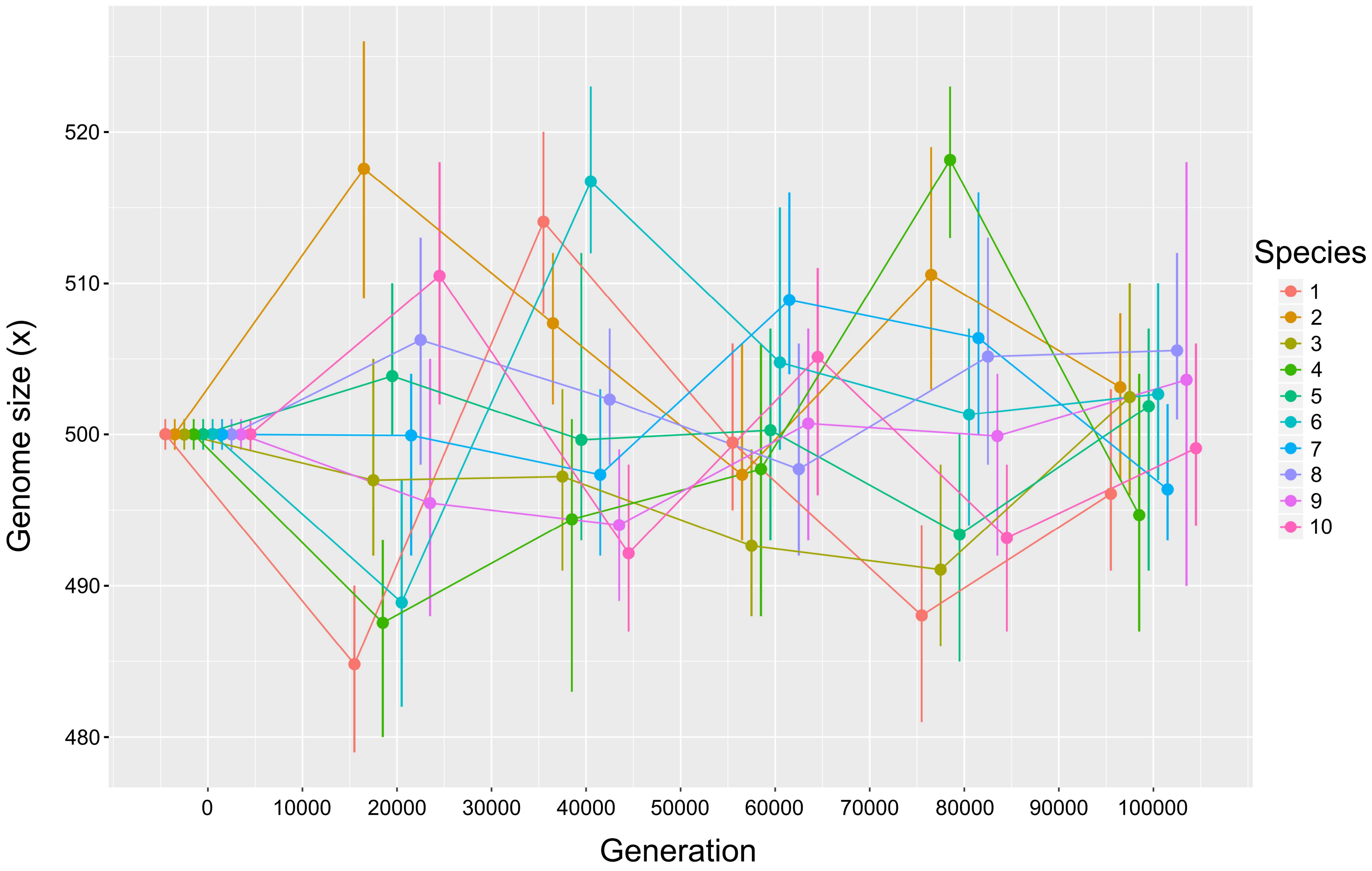

Genome size (x)
520
510
500
490
480
0
10000
20000
30000
40000
50000
60000
70000
80000
90000
100000
Generation
Species
1
2
3
4
5
6
7
8
9
10

M

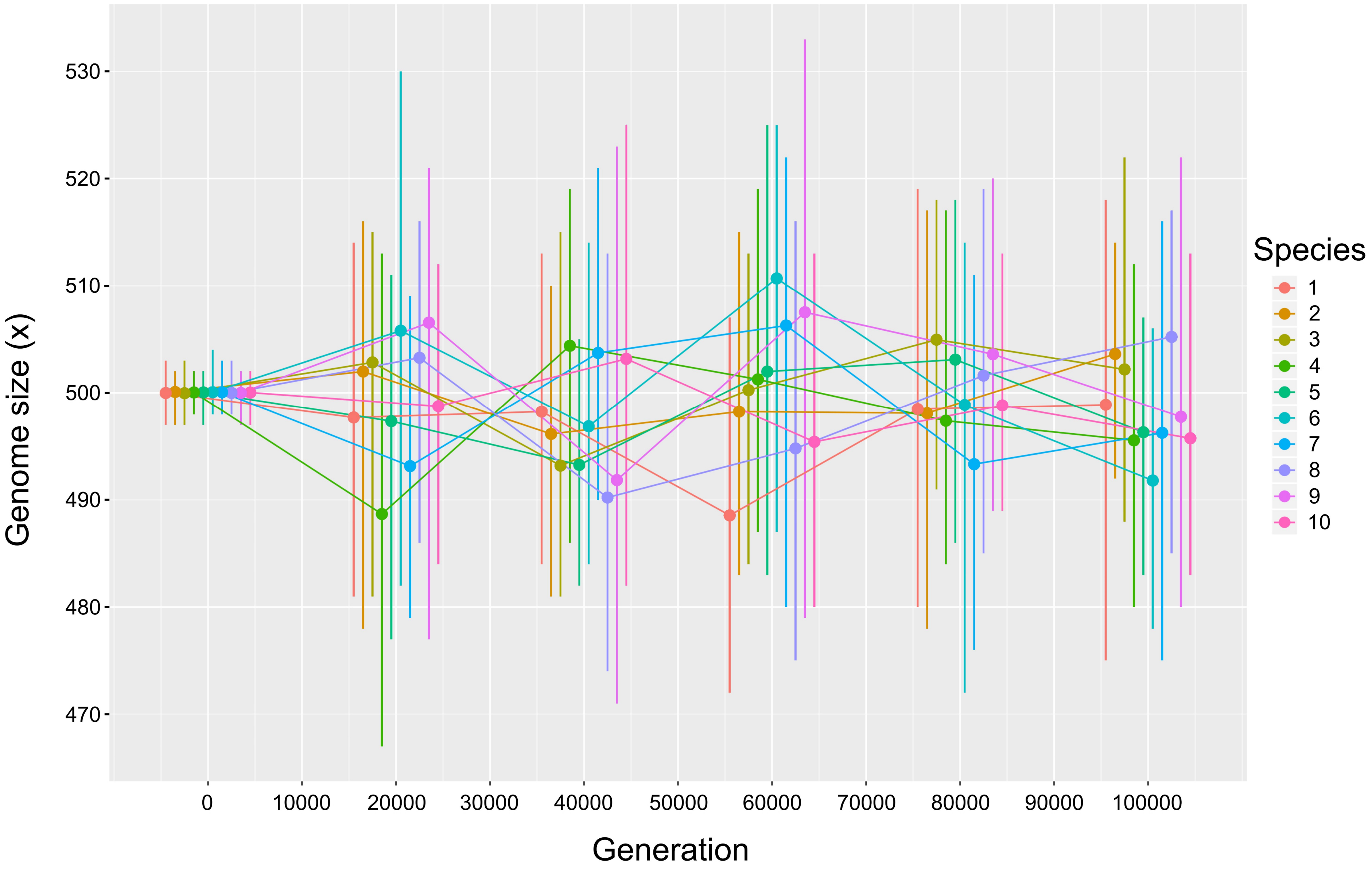

530
520
510
500
490
480
470
Genome size (x)
0
10000
20000
30000
40000
50000
60000
70000
80000
90000
100000
Generation
Species
1
2
3
4
5
6
7
8
9
10

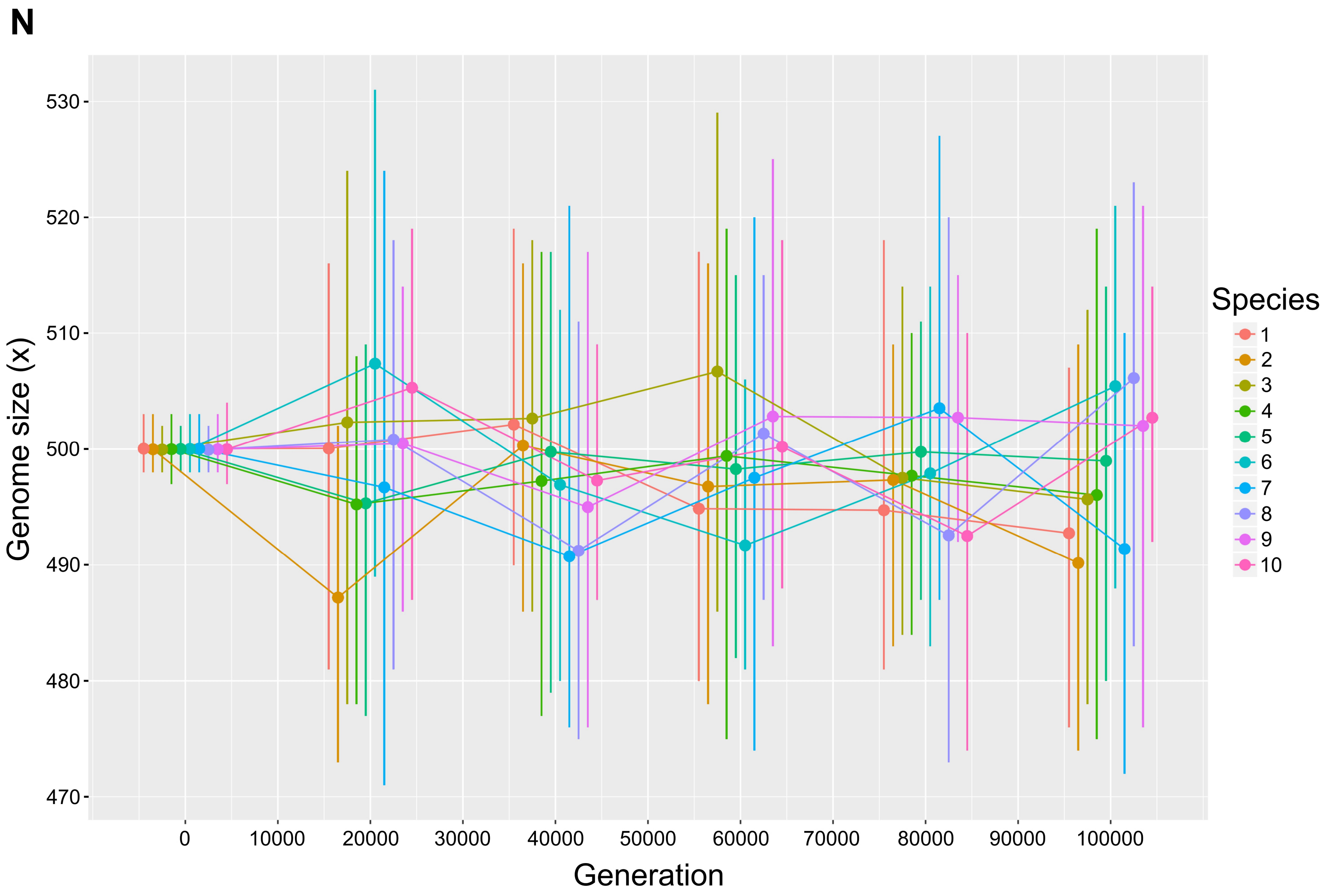
N
Genome size (x)
530
520
510
500
490
480
470
0
10000
20000
30000
40000
50000
60000
70000
80000
90000
100000
Generation
Species
1
2
3
4
5
6
7
8
9
10

O

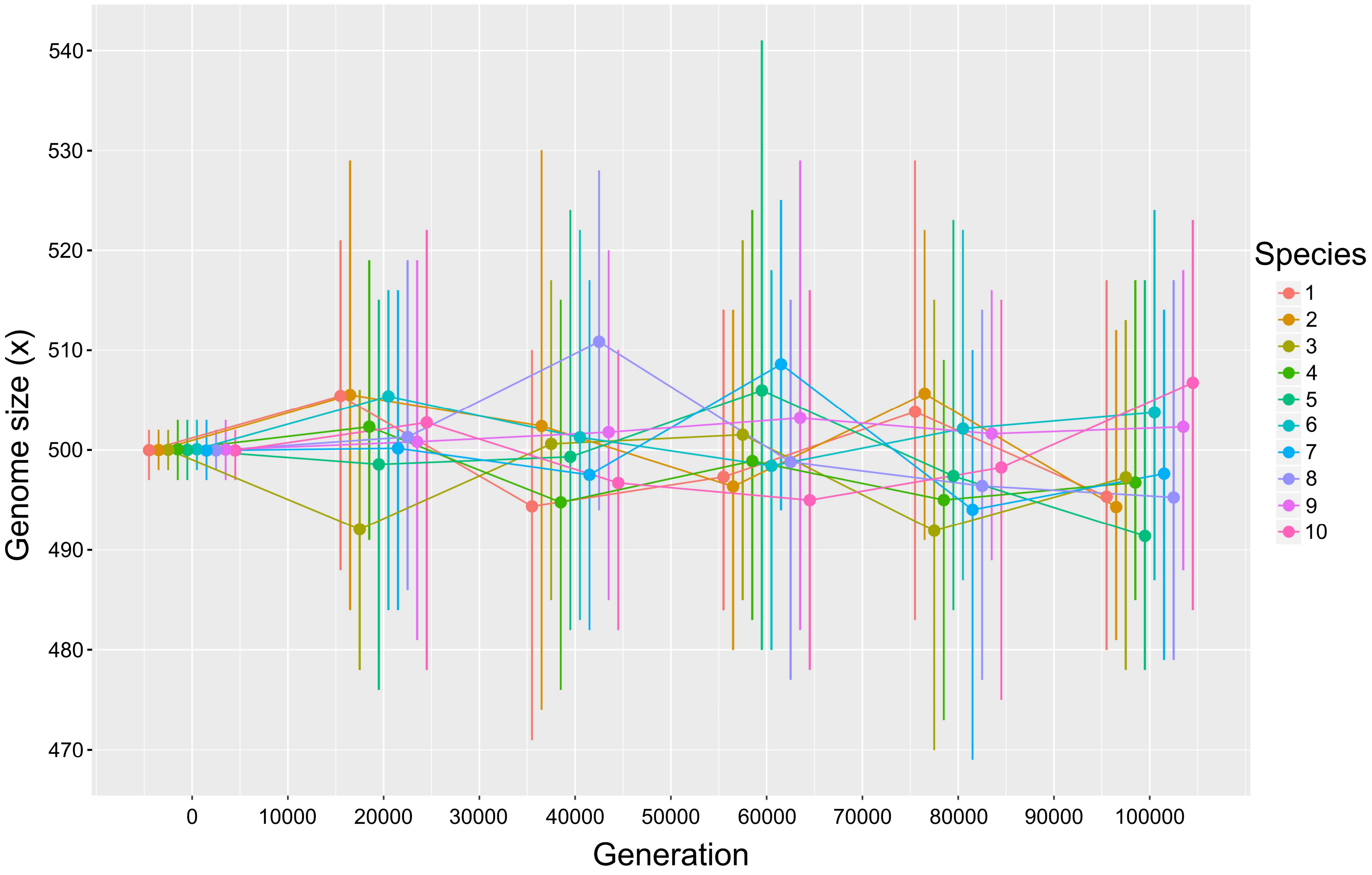

Genome size (x)
540
530
520
510
500
490
480
470
0
10000
20000
30000
40000
50000
60000
70000
80000
90000
100000
Generation
Species
1
2
3
4
5
6
7
8
9
10

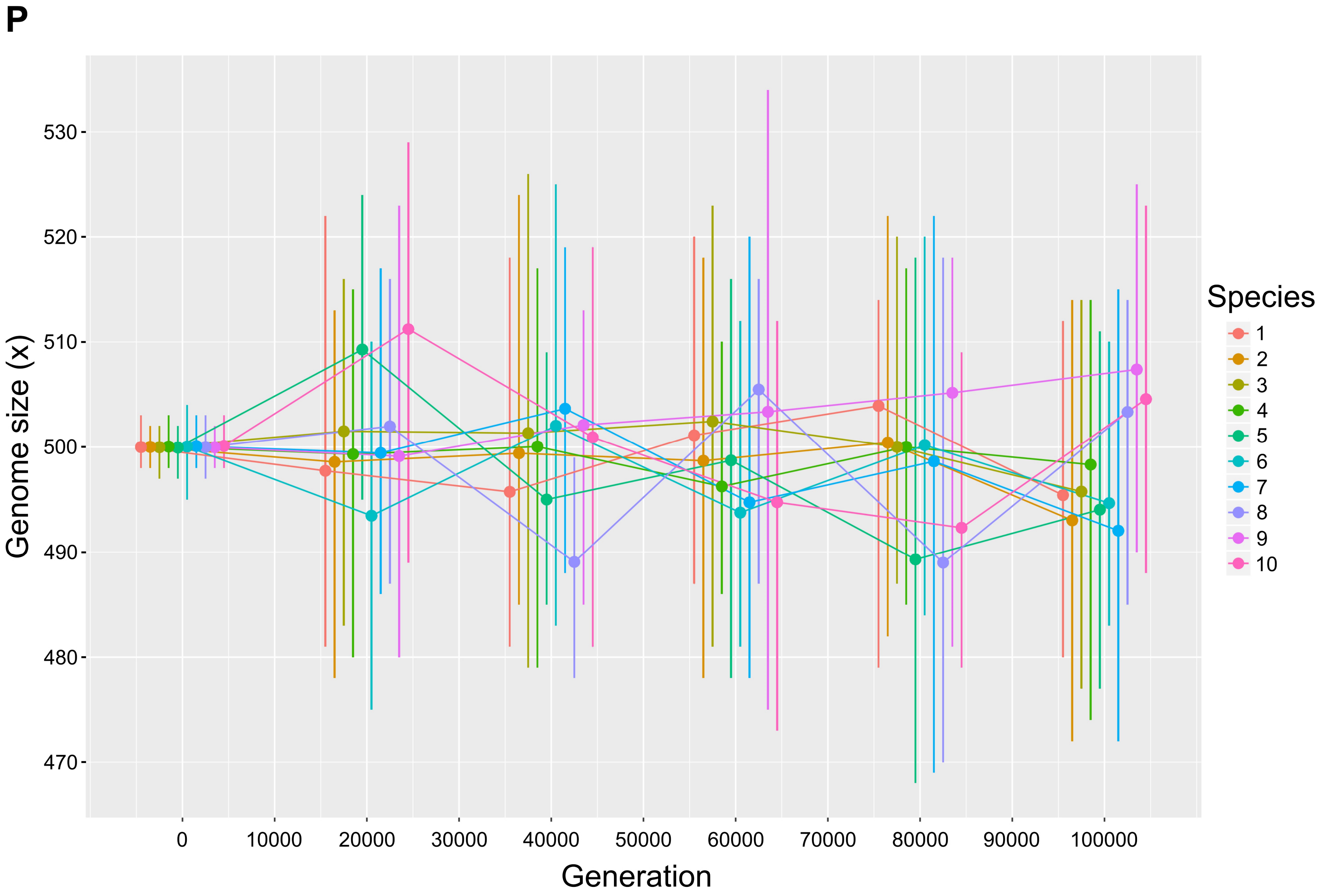
P
Genome size (x)
530
520
510
500
490
480
470
Generation
0
10000
20000
30000
40000
50000
60000
70000
80000
90000
100000
Species
1
2
3
4
5
6
7
8
9
10
